# Dimer opening enables brain-type creatine kinase to sense membrane curvature

**DOI:** 10.64898/2026.08.04.742861

**Authors:** Samantha Gies, Nolan K. McLaughlin, Orianna H. Kou, Ahmed Shubbar, Chen Kong, Tianqi Wu, Sana Khan, Alessandro Ustione, Ian Miller, David W. Piston, Mahmoud Moradi, Michael L. Gross, Wade F. Zeno, Reza Dastvan

## Abstract

Brain-type creatine kinase (CK-BB) buffers local ATP demand through reversible phosphotransfer between ATP and phosphocreatine, yet how this soluble metabolic enzyme engages membrane compartments is unknown. Here, we combine fluorescence microscopy, DEER spectroscopy, hydrogen–deuterium exchange and native mass spectrometry, DEER-and AlphaFold-guided modeling, and long-timescale molecular dynamics to define the pH- and substrate-regulated conformational landscape governing CK-BB membrane association. Acidification promotes curvature-sensitive membrane binding and redistributes endogenous and recombinant CK-BB from diffuse cytosolic pools to punctate vesicular structures and membrane ruffles. Substrates independently promote curvature-sensitive association at neutral pH. DEER and modeling reveal an asymmetric dimer in which the convex surface remains restrained, whereas the concave catalytic–regulatory surface samples pH- and substrate-dependent intermediates. We identify progressive dimer opening as a novel regulatory mechanism whereby acidification and substrate binding increase dynamics across the convex surface and N-terminal dimer interface, generating membrane-competent conformations that facilitate curvature sensing and membrane association. Substrate binding buffers acid-induced deprotection while preserving dynamics near the His191/Ser199 regulatory interface. These findings establish CK-BB as a previously unrecognized curvature-sensitive metabolic enzyme and define dimer dynamics as a molecular switch coupling protonation and substrate occupancy to curved-membrane recognition and localized ATP regeneration, with potential relevance to neurodegeneration and cellular stress.

## Introduction

Creatine kinases (CKs) form a phosphagen circuit that buffers ATP and ADP in tissues and subcellular compartments with high and fluctuating energy demand^1–6^. Cytosolic brain-type creatine kinase (CK-BB) functions as a homodimer that catalyzes reversible phosphoryl transfer between ATP and creatine (Cr), producing ADP, H^+^ and phosphocreatine (PCr). This reaction supports rapid ATP regeneration at sites of localized energy consumption, including neuronal processes, ion-transport microdomains, actomyosin assemblies, cortical membrane compartments, and other CK-BB-enriched subcellular microdomains^1,4–10^.

CK-BB has emerged as a regulator of cell morphology, motility and metastatic progression. Its cellular distribution is altered in metastatic cancer cells^3,11–13^, and CK-BB has been detected extracellularly or as a serum-associated marker in several cancers, including colorectal, ovarian, neuroblastoma, prostatic and lung cancer^11,12,14,15^. CK-BB-dependent phosphagen metabolism has also been linked to metastatic survival, hypoxia, invasion and therapeutic vulnerability^11,13,16–23^. Yet, despite its extracellular detection and functional association with metastatic energetics^11,19^, CK-BB lacks a canonical secretion signal peptide. These observations raise the possibility that CK-BB engages mechanisms relevant to unconventional protein secretion (UPS), in which leaderless cytosolic proteins bypass the endoplasmic reticulum and Golgi apparatus through pathways that often involve lipid binding, membrane insertion, or trafficking through endosomal and secretory intermediates, particularly under cellular stress^24,25^. Consistent with this framework, CK-BB transiently accumulates in actin-rich membrane ruffles and phagocytic cups, and loss or inhibition of CK-BB activity impairs cell spreading, migration and complement-mediated phagocytosis^9,10,26^. Together, these findings suggest that CK-BB function depends not only on catalytic chemistry, but also on regulated localization, stress-dependent membrane access, and conformational adaptability.

Prior biochemical observations further support the potential for CK-BB membrane engagement. CK isoforms interact with phospholipid monolayers^27^, and CK-BB or CK activity has been associated with synaptosomal membranes and synaptic vesicles^6,28,29^. In Alzheimer’s disease brain and oxidative-stress models, CK-BB modifications reduce activity, impair nucleotide binding, and coincide with aberrant membrane partitioning or focal creatine deposits^30–32^. Together, these findings suggest that CK-BB disease-associated functions may depend on controlled access to membrane-proximal compartments as well as catalytic regulation. However, whether CK-BB directly engages defined membrane surfaces in response to environmental or metabolic cues, and whether this interaction is modulated by membrane curvature, remain unclear.

Acidification is particularly relevant to CK-BB biology. Hypoxic and metabolically stressed metastatic cancer cells experience altered intracellular and extracellular pH, and acidic microenvironments promote metastasis, invasion, secretion, vesicle trafficking and cytoskeletal remodeling in cancer and inflammatory contexts^33–36^. Disrupted intracellular pH homeostasis is also associated with neuroinflammatory and neurodegenerative states, including neuronal cytosolic acidification and excessive endosomal acidification in AD-relevant astrocytes^37,38^. In parallel, endosomal and secretory compartments maintain regulated pH gradients that can reshape protein–membrane interactions. Acute extracellular acidification therefore provides a tractable test of whether an acid-responsive CK-BB state can redistribute the enzyme from a diffuse cytosolic pool to vesicular or cortical membrane compartments. For a soluble enzyme enriched at membrane ruffles and vesicular structures, key questions are whether acidification or substrate availability remodels the CK-BB dimer into membrane-competent conformations and whether membrane curvature modulates their engagement.

High-resolution structures of human and chicken CK-BB have captured the dimeric ligand-free protein^39,40^ and ligand-bound states^41^, showing that nucleotide and transition-state analogue binding promotes active-site closure and rearranges catalytic loops required for substrate recognition and phosphoryl transfer^41^. Complementary structural and biochemical studies have identified catalytically important flexible loops, active-site histidines, nucleotide-binding arginine residues, and loop movements that occlude the active site during catalysis^42–45^. However, the structural and dynamic basis by which CK-BB adapts to acidification, substrate binding, and membrane-proximal environments remains poorly understood. This gap is important because CK-BB operates in settings where pH, substrate availability, ionic composition, and membrane curvature can vary locally, including hypoxic tumors, secretory and vesicular compartments, synaptic membranes, and actin-rich cortical domains.

Here, we integrate membrane biophysics with structural dynamics to define how acidic pH and ligand binding remodel CK-BB. Immunofluorescence shows that acidification redistributes CK-BB to vesicular and membrane-ruffle compartments^26,46^, while quantitative tethered-vesicle fluorescence microscopy demonstrates direct, curvature-sensitive membrane association that is enhanced by acidic pH and, independently, by substrate availability at neutral pH. Double electron-electron resonance spectroscopy (DEER/PELDOR)^47–52^ maps inter-protomer conformational equilibria across the convex and concave dimer surfaces, and DEER- and AlphaFold-guided modeling^53–56^ defines the underlying conformational landscape and its modulation by protonation and ligand binding. Hydrogen–deuterium exchange mass spectrometry (HDX-MS)^57–62^ provides comprehensive peptide-level readouts of local backbone dynamics and solvent accessibility, while molecular dynamics simulations^63–67^ connect these exchange signatures to structural motion. Together, these approaches reveal a pH- and ligand-gated CK-BB energy landscape in which dimer opening and selective remodeling of the convex and concave surfaces generate membrane-competent states, providing a structural mechanism that couples local ATP regeneration to curved vesicular and actin-remodeling membranes and identifies CK-BB as a previously unrecognized curvature-sensitive member of the creatine kinase family.

## Results

### Functional and structural integrity of CK-BB constructs

For DEER measurements and fluorescence microscopy, single-cysteine substitutions were introduced into a cysteine-less (CL) CK-BB background to enable site-specific spin labeling. The five native cysteines were replaced by residues found in homologs or by substitutions previously shown to preserve activity and stability, yielding CL CK-BB^68,69^. Although the conserved Cys283, likely through its deprotonated thiolate state, contributes to maximal creatine kinase activity, current evidence does not support a direct catalytic role or a role in substrate-induced conformational changes^70^. We assessed the functional integrity of the CL construct relative to wild-type (WT) CK-BB using a creatine kinase activity assay that quantifies CK-dependent ATP generation from phosphocreatine and ADP. Mutation of Cys283 to serine strongly reduced activity, whereas the C283D-containing CL background retained measurable activity and was therefore selected for DEER and fluorescence microscopy analyses (Supplementary Fig. 1a). Importantly, restoring Cys283 in the CL background, as in the CL^D283C^ construct, showed that mutation of the remaining native cysteines did not impair catalytic activity. Consistent with the pH-dependent reversibility of the CK reaction, WT CK-BB activity in the ATP-generating direction was higher at mildly acidic pH than at pH 7.3 (Supplementary Fig. 1b), supporting the functional relevance of the pH range used for structural-dynamics measurements^70^. Far-UV circular dichroism spectra of WT, CL and spin-labeled mutants were closely similar, confirming that cysteine substitutions and spin labeling did not perturb global secondary structure (Supplementary Fig. 1c). The complementary HDX-MS experiments were performed with WT protein.

### Acidification drives curvature-sensitive membrane recruitment of CK-BB

Because CK-BB is recruited to actin-rich cortical microdomains, supports local ATP generation during actin remodeling, and is implicated in extracellular metabolic energetics, we first tested whether acidification changes the cellular distribution of CK-BB. HCT116 colorectal carcinoma and HEK293 cells expressing endogenous CK-BB or recombinant FLAG-tagged CK-BB were incubated at neutral pH and pH 5.5 and analyzed by immunofluorescence (Fig. 1). At pH 7.2, CK-BB was largely diffuse and cytosolic in both cell types, consistent with its canonical role as a soluble phosphotransfer enzyme (Fig. 1a). In contrast, acidification to pH 5.5 increased punctate and vesicular staining and enriched CK-BB at membrane ruffles (Fig. 1b), a pattern observed for endogenous CK-BB and for recombinant FLAG-CK-BB detected with either anti-CK-BB or anti-FLAG antibodies. These cellular data extend prior observations that CK-BB accumulates in phagocytic cups, membrane ruffles, and other actin-rich protrusions^9,10,26^.

**Figure 1.**
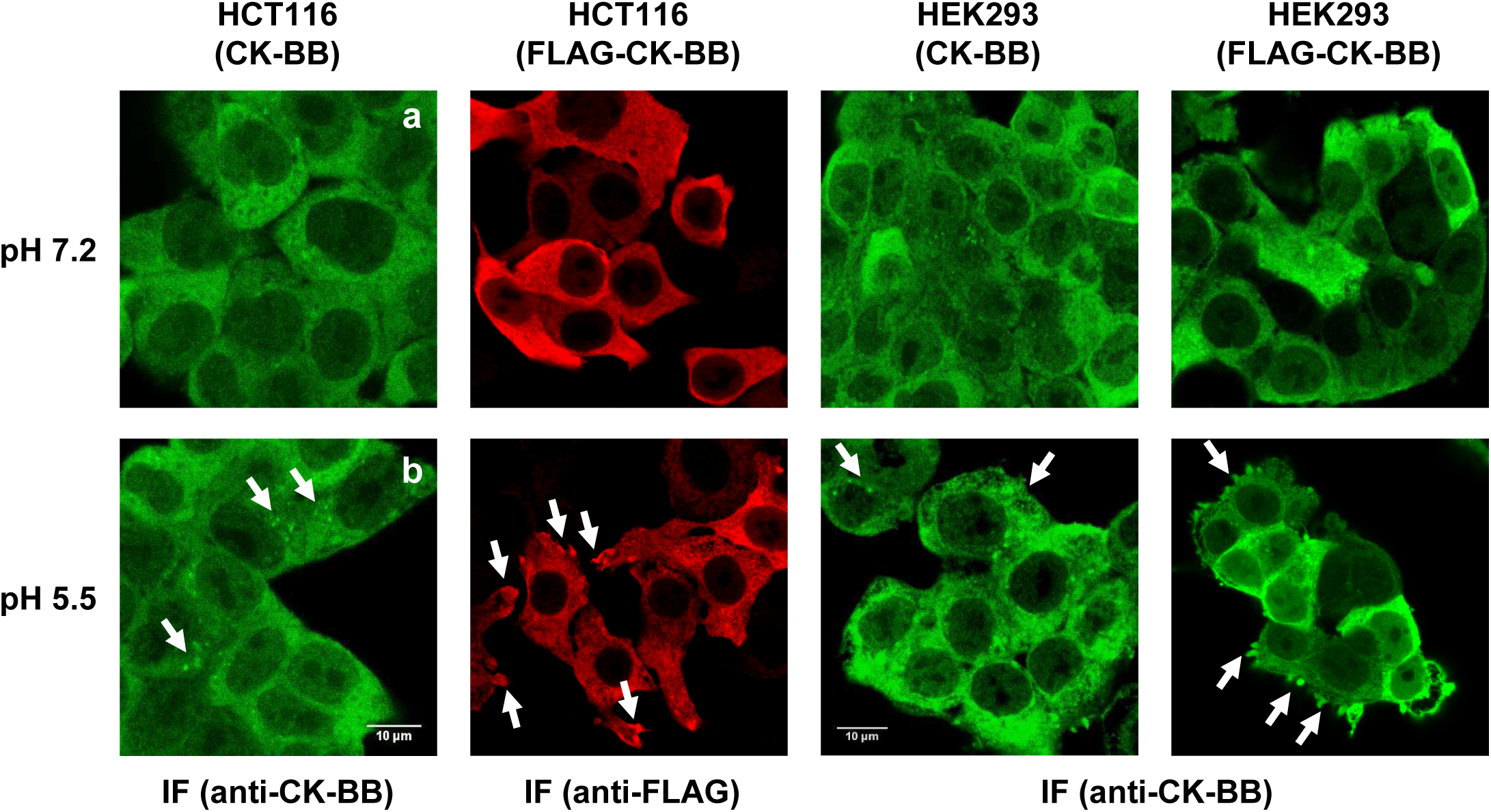
Acidic conditions redistribute CK-BB to vesicles and membrane ruffles. Immunofluorescence staining of HCT116 colorectal carcinoma and HEK293 cells expressing endogenous CK-BB or recombinant FLAG-tagged CK-BB after incubation at pH 7.2 (**a**) or pH 5.5 (**b**) in a 5% CO_2_ incubator. Endogenous CK-BB was detected with an anti-CK-BB antibody (green), while recombinant FLAG-CK-BB was detected with anti-FLAG (red) or anti-CK-BB antibodies, as indicated. Compared with neutral pH, acidic conditions increased punctate/vesicular staining and enriched CK-BB at membrane ruffles. Scale bars, 10 μm.

To determine whether acidification-induced CK-BB redistribution reflects direct membrane association, we used a tethered-vesicle fluorescence assay (Fig. 2a)^71,72^. ATTO 647N-DPPE was incorporated into lipid vesicles, and its fluorescence intensity was calibrated against dynamic light scattering (DLS) measurements to determine vesicle diameter (Fig. 2b). CK-BB binding was monitored using an AZDye-488-labeled N27C variant, enabling single-molecule quantification of membrane-bound CK-BB molecules on each vesicle (Fig. 2c). Plasma membrane (PM)-mimetic vesicles containing PC, PE, PS, PI, and cholesterol showed no detectable CK-BB association at pH 7.3, whereas vesicle-bound CK-BB was readily apparent at pH 5.5 (Fig. 2d). Quantification confirmed no measurable binding at pH 7.3 at concentrations up to 500 nM. By contrast, at pH 5.5, binding was detectable at 100 nM and increased progressively at 250 and 500 nM (Fig. 2e). Strikingly, analysis of membrane-bound protein density revealed preferential association with smaller vesicles, with CK-BB density increasing sharply as vesicle diameter decreased (Fig. 2f). Normalized density profiles showed at least tenfold enrichment on the smallest vesicles at all three CK-BB concentrations, revealing pronounced curvature-sensitive membrane association under acidic conditions (Fig. 2g). This curvature sensitivity was even stronger on synaptic vesicle (SV)-mimetic membranes composed of PC, PE, PS and cholesterol, approaching 40-fold enrichment on the smallest vesicles (Supplementary Fig. 2). Thus, acidification reveals a previously unrecognized capacity of CK-BB to preferentially engage highly curved membranes, establishing CK-BB as a curvature-sensitive member of the creatine kinase family.

**Figure 2.**
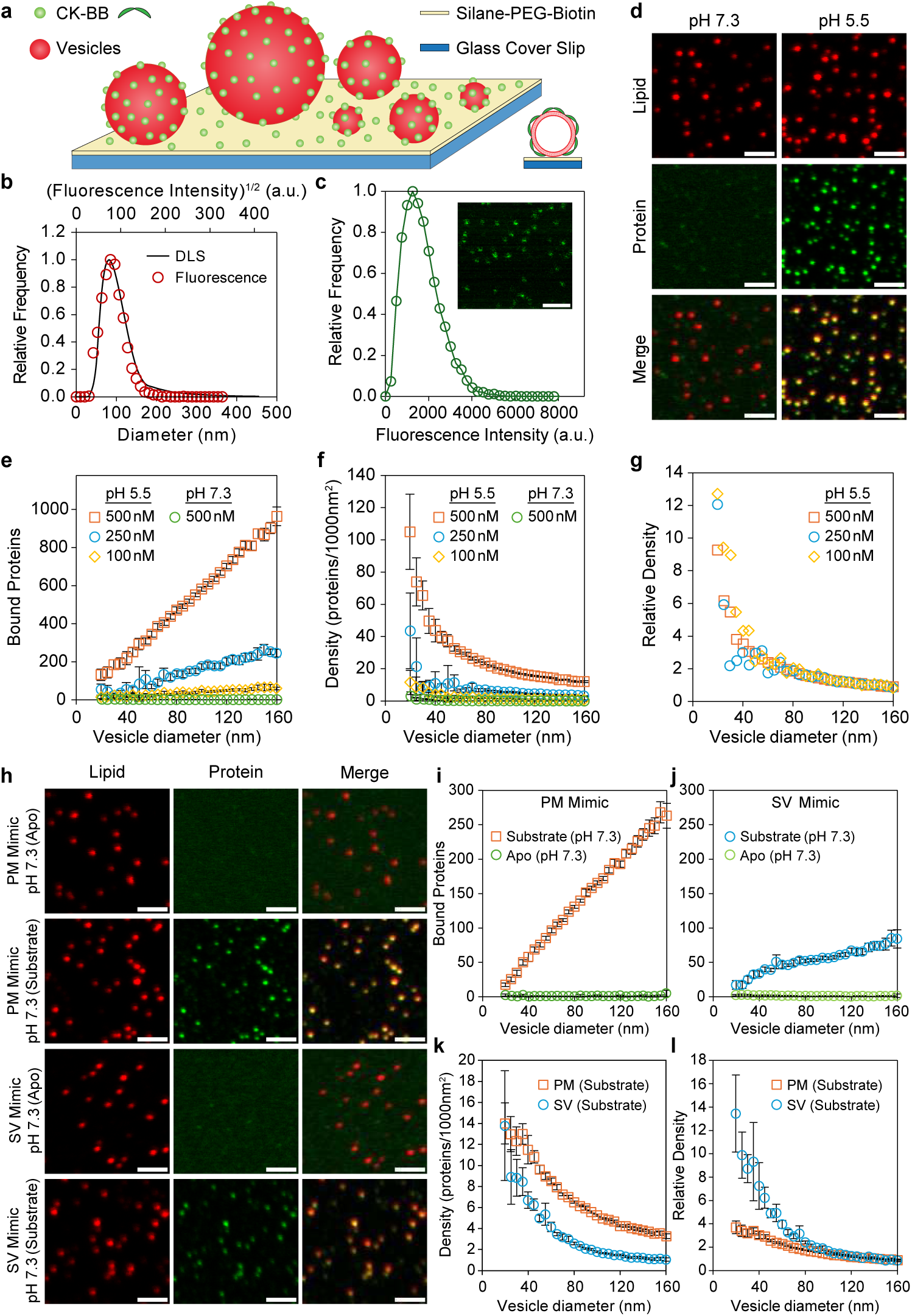
Acidification and substrate binding independently promote curvature-sensitive CK-BB membrane association. (**a**) Schematic of the tethered-vesicle assay. (**b**) Representative lipid-fluorescence calibration used to determine vesicle diameter. (**c**) Representative single-molecule CK-BB fluorescence calibration used to quantify membrane-bound protein. (**d**) Lipid, CK-BB and merged fluorescence micrographs of tethered plasma membrane (PM)-mimetic vesicles incubated with apo CK-BB at pH 7.3 or 5.5. (**e**) Number of membrane-bound apo CK-BB molecules as a function of vesicle diameter. (**f**) Membrane-bound CK-BB density as a function of vesicle diameter. (**g**) Relative CK-BB density normalized to the mean density on vesicles 130–160 nm in diameter. (**h**) Lipid, CK-BB and merged fluorescence micrographs of PM- and synaptic vesicle (SV)-mimetic vesicles incubated with apo or substrate-bound CK-BB at pH 7.3. (**i**,**j**) Number of membrane-bound CK-BB molecules as a function of vesicle diameter for PM (**i**) and SV (**j**) mimetics. (**k**) Membrane-bound CK-BB density as a function of vesicle diameter for PM and SV mimetics. (**I**) Relative CK-BB density normalized to that on 130–160-nm vesicles. Substrate-dependent binding is greater on PM mimetics, whereas the relative enrichment on highly curved vesicles is more pronounced for SV mimetics. For **e**–**g** and **i**–**I**, vesicles 20–160 nm in diameter were grouped into 29 bins at 5-nm intervals, and each point represents the mean within the corresponding bin. Numbers of imaged and processed vesicles were *N* = 2554 (pH 5.5, 100 nM), 1927 (pH 5.5, 250 nM), 12,006 (pH 5.5, 500 nM), 3261 (pH 7.3, 500 nM), 3825 (SV mimic, apo), 11,554 (SV mimic, substrates), 4497 (PM mimic, apo) and 12,039 (PM mimic, substrates). Data for apo CK-BB at pH 5.5, 500 nM and all substrate-bound conditions were pooled across three biological replicates (*n* = 3); all other conditions represent individual measurements (*n* = 1). Error bars indicate 95% confidence intervals of the mean. Scale bars, 2 μm.

These findings also align with the UPS-related premise that stress-associated acidification, vesicular trafficking, membrane curvature and actin remodeling can create routes for leaderless cytosolic proteins to access membrane-proximal compartments^24,25^. Together with the tethered-vesicle measurements, they show that acidification not only redistributes CK-BB toward vesicular and cortical membrane compartments in cells, but is also sufficient to drive its direct, curvature-sensitive association with model membranes. Although these measurements do not establish secretion or membrane translocation, they define an acid-responsive, curvature-sensitive membrane-binding state for CK-BB, with potential relevance to CK-BB dysfunction in neurodegeneration and cellular stress^30–32^.

### Substrate binding promotes curvature-sensitive membrane association at neutral pH

Because CK-BB physiologically operates in the presence of nucleotide and phosphagen substrates, we examined whether substrate binding promotes membrane association independently of acidification. Using the tethered-vesicle assay, we measured CK-BB binding at pH 7.3 in the presence of 10 mM MgATP, 0.8 mM ADP, 20 mM PCr and 10 mM Cr, approximating a substrate-rich phosphagen-buffering state and favoring ligand-occupied CK-BB conformations^73^ (Fig. 2h–l). Whereas apo CK-BB showed no measurable membrane association at pH 7.3 even at 1000 nM, substrates promoted detectable binding. Notably, membrane-bound CK-BB was preferentially enriched on smaller vesicles, showing that substrate-driven recruitment remains curvature-sensitive. The magnitude of binding under substrate-bound conditions at pH 7.3 was nevertheless comparable to or lower than that of apo CK-BB at pH 5.5, despite the higher protein concentration required (Fig. 2 and Supplementary Fig. 2). Thus, substrate binding independently promotes curvature-sensitive CK-BB membrane association at neutral pH, although acidification elicits a stronger response under these conditions.

In addition, we used fluorescence recovery after photobleaching (FRAP) to probe the dynamics of membrane-associated CK-BB across membrane compositions and solution conditions (Supplementary Fig. 3). Fluorescence recovery remained ≤10% over the four-minute measurement window, indicating that most membrane-associated CK-BB forms a persistent, slowly exchanging population. Neither the extent of recovery nor the apparent exchange kinetics varied systematically with vesicle diameter. Thus, the pronounced curvature dependence observed at steady state primarily reflects preferential CK-BB accumulation on highly curved membranes rather than curvature-dependent differences in membrane exchange dynamics.

Together, these measurements establish protonation and substrate binding as independent physiological inputs promoting CK-BB engagement with curved membranes. These findings provide an experimental foundation for defining how protonation and ligand binding remodel the CK-BB dimer to generate membrane-competent conformational states. The persistence of curvature sensitivity under two distinct regulatory conditions further indicates that membrane-curvature sensing is an intrinsic property of membrane-competent CK-BB states rather than a consequence of acidification alone.

### The convex and concave faces of CK-BB sample distinct conformational equilibria

To define how pH and substrate binding reshape the CK-BB dimer, we placed inter-protomer spin-label reporters across two major surfaces: the convex face, which spans the NTD dimer interface and C-terminal β-sheet core, and the concave face, which contains active-site and regulatory elements (Figs. 3 and 4). On the convex surface, most reporter pairs showed narrow distance distributions and limited redistribution between pH 7.3 and 6.5 in both apo and ligand-bound states (Fig. 3 and Supplementary Fig. 4). More pronounced changes were observed primarily at pH 5.5, indicating that the convex surface is comparatively restrained under neutral and mildly acidic conditions but becomes more responsive upon stronger acidification.

**Figure 3.**
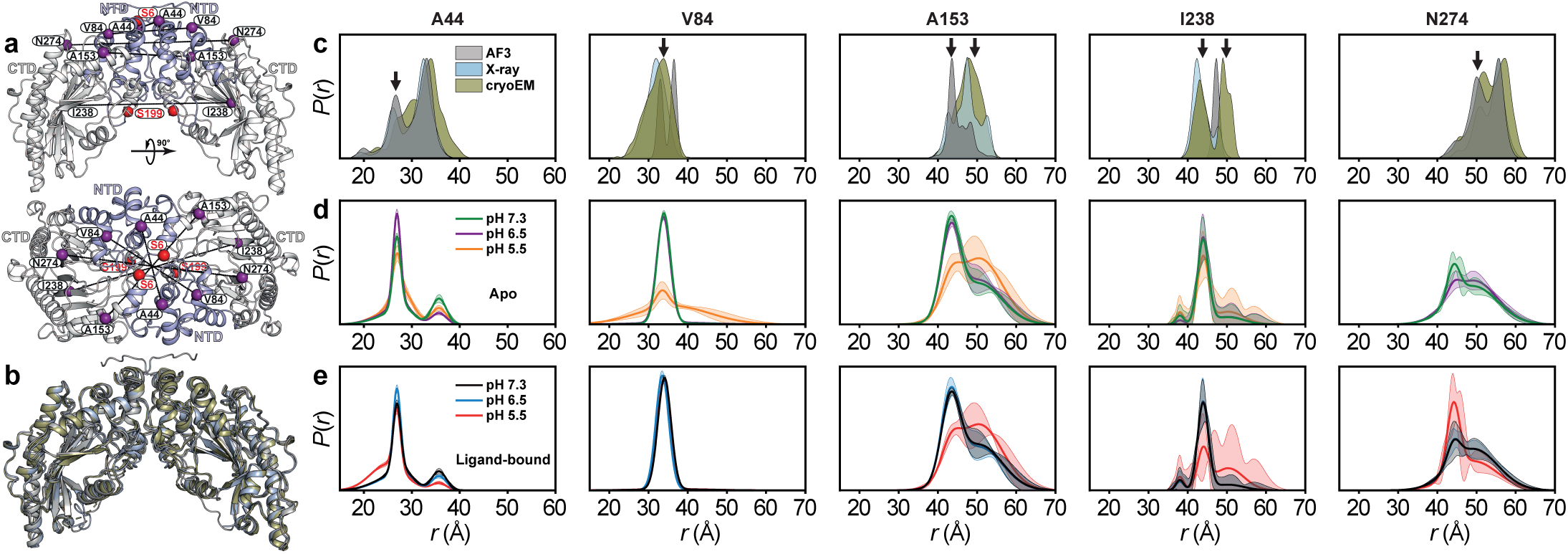
Conformational dynamics of the CK-BB dimer at the convex side of the NTD interface and CTD beta-sheet core. (**a**) Inter-protomer spin-label pairs (purple spheres) used for DEER measurements across the convex side of the NTD dimer interface and the C-terminal beta-sheet core. Putative phosphorylation sites Ser6 and Ser199 are shown as red spheres; NTD and CTD regions are colored light blue and grey, respectively. (**b**) Overlay of cryo-EM (green; PDB 6V9H), X-ray (blue; PDB 7TUN) and AlphaFold (grey) CK-BB structures. (**c**) Predicted spin-label distance distributions from the structures, with arrows marking components corresponding to experimentally observed DEER intermediates. (**d**,**e**) DEER distance distributions, *P*(*r*), for apo (**d**) and ligand-bound (**e**) CK-BB at physiological and acidic pH. Except under strongly acidic conditions (pH 5.5), these CK-BB regions remain relatively rigid and show minimal pH- or ligand-dependent changes. Confidence bands indicate 2σ uncertainty from fitting of the primary DEER traces. Raw DEER decays and fits are shown in Supplementary Fig. 4.

**Figure 4.**
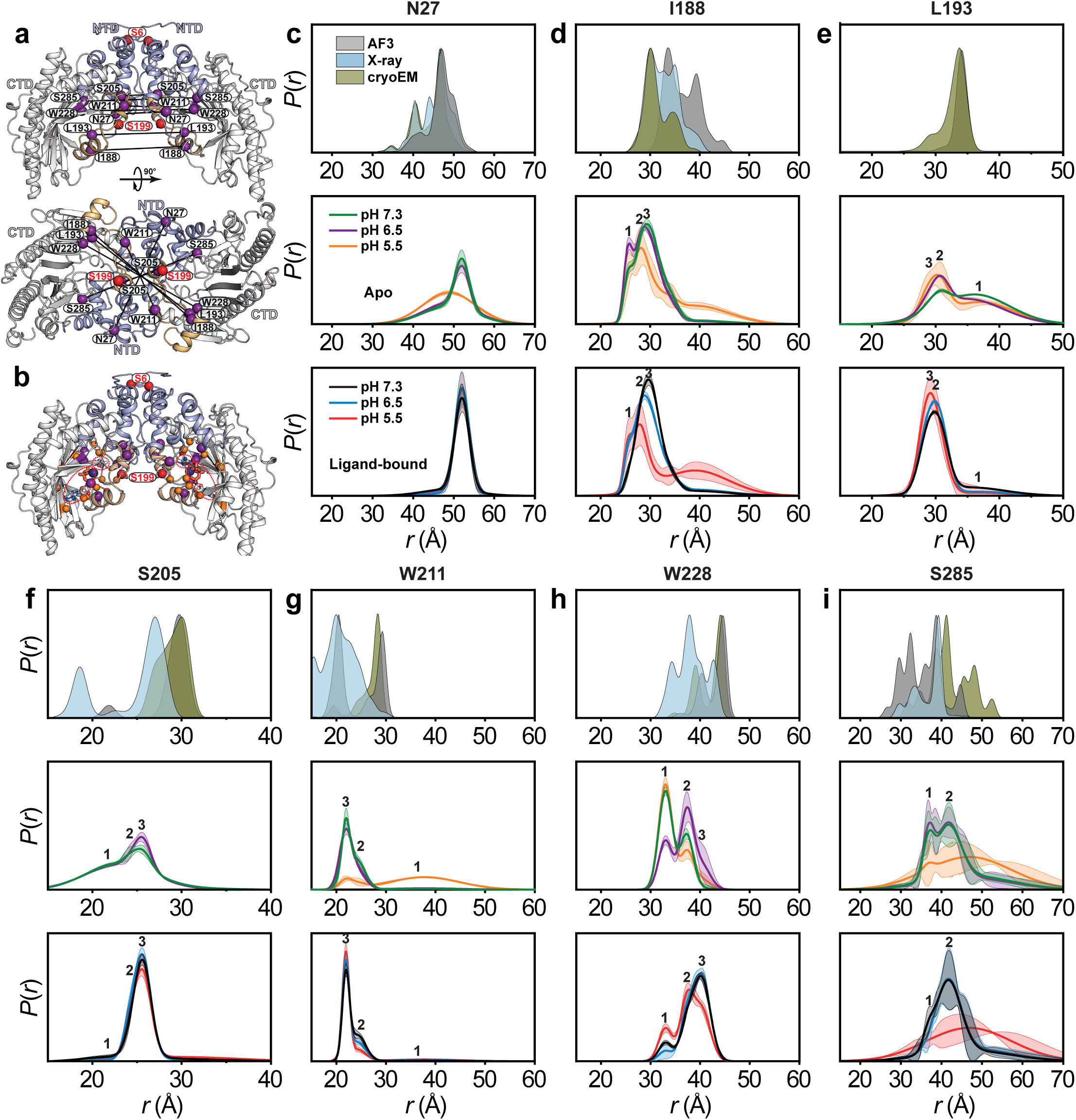
The dynamic concave side of the CK-BB dimer undergoes pH- and ligand-dependent conformational remodeling. (**a**) Inter-protomer spin-label pairs (purple spheres) used for DEER measurements across the concave side of the CK-BB dimer. Ser6 and Ser199 phosphorylation sites are shown as red spheres; NTD and CTD regions are colored light blue and grey, respectively, and the region flanking Ser199 is highlighted in light orange. (**b**) AlphaFold3 model of CK-BB bound to Mg^2+^ATP and creatine, with catalytic and substrate-binding residues shown as orange spheres. (**c**–**i**) Predicted spin-label distance distributions from AF3, X-ray and cryo-EM structures (top) are compared with DEER-derived distance distributions, *P*(*r*), for apo (middle) and ligand-bound CK-BB (bottom) at physiological and acidic pH. The concave side, particularly the Ser199-containing region, is highly dynamic, displays pH- and ligand-dependent conformational changes, and samples three distinct conformational intermediates. Confidence bands indicate 2σ uncertainty from fitting of the primary DEER traces. Raw DEER decays and fits are shown in Supplementary Fig. 5.

The concave face displayed a markedly different behavior. Reporters surrounding the Ser199-containing region and neighboring active-site elements sampled multiple conformational intermediates that redistributed with both pH and ligand binding (Fig. 4 and Supplementary Fig. 5). Several reporter pairs resolved three distance populations: two relatively narrow, well-defined components and a broader, more dynamic population that became more prominent under acidic conditions, particularly at pH 5.5. Although some experimental components overlapped with distances predicted from AlphaFold3, X-ray, and cryo-EM structures, the DEER data revealed broader ensembles and intermediate states that were not captured by any single model.

Together, these results show that CK-BB is dynamically polarized. The convex face remains comparatively constrained across most conditions, whereas the concave face, particularly the region containing the nucleotide-binding residue His191 and the putative Ser199 phosphorylation site, is strongly coupled to protonation and ligand occupancy. This face-specific remodeling links CK-BB conformational equilibria to conditions that promote membrane association.

### Protonation and ligand binding reveal dimer opening as a regulatory mechanism

We next translated the DEER-derived distance constraints into structural ensembles using DEERFold^53^, a fine-tuned AlphaFold2-based network^55^ that incorporates DEER distance distributions as distogram inputs to predict spin-label-constrained conformations. Single-Gaussian approximations of the stabilized distance components observed under each condition were used as DEERFold restraints. Principal component analysis resolved two dominant collective motions of the CK-BB dimer (Fig. 5a). PC1 described lateral dimer expansion, whereas PC2 captured hinge-like opening and closure, as illustrated by the corresponding morphing trajectories (Supplementary Videos 1 and 2). Within this conformational space, ligand-bound CK-BB at pH 5.5 shifted most strongly toward the laterally expanded, open ensemble. Ligand-bound CK-BB at pH 7.3 occupied an intermediate region, whereas apo ensembles remained comparatively closed (Fig. 5a). Independently, we refined the unbiased AlphaFold3 (AF3) model^56^ with the same DEER constraints, yielding similar conformations (Fig. 5b and Supplementary Figs. 6 and 7)^54^.

**Figure 5.**
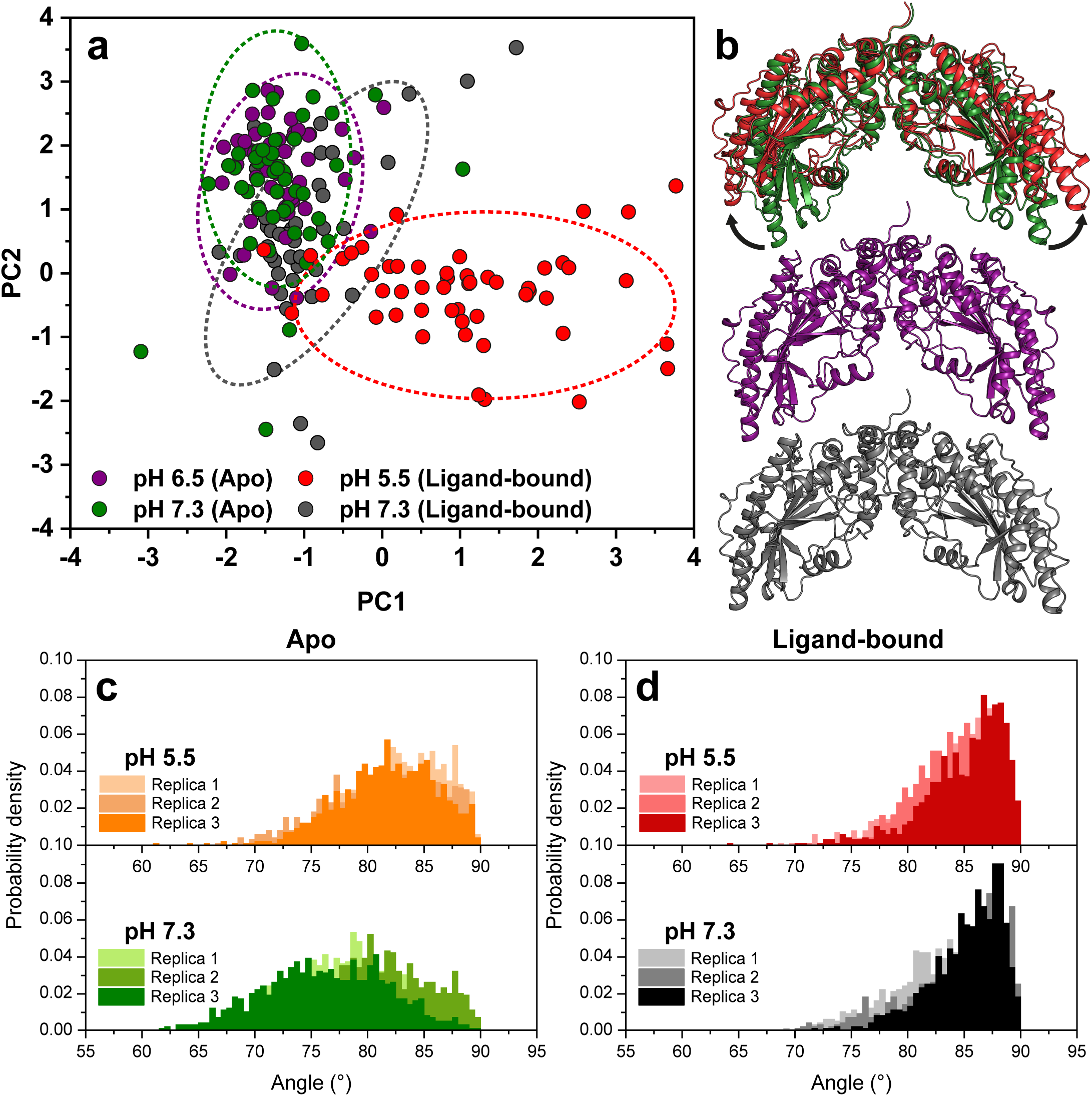
Acidic pH and ligand binding promote opening of the CK-BB dimer. (**a**) Principal component analysis of DEER-guided AlphaFold models. PC1 captures lateral dimer expansion, whereas PC2 reports hinge-like opening and closure of the CK-BB dimer (Supplementary Videos 1 and 2). Ligand-bound CK-BB at pH 5.5 occupies the most expanded and open conformational space, while ligand-bound CK-BB at pH 7.3 shows an intermediate distribution and apo states remain comparatively closed. (**b**) DEER-refined AF3 models of apo and ligand-bound CK-BB under physiological and acidic conditions (Supplementary Figs. 6 and 7), showing a transition from a more closed, dynamic apo ensemble at pH 7.3 to a more open ligand-bound conformation under acidic conditions. (**c**,**d**) Dimer-opening angle distributions from triplicate 1-µs MD simulations of apo (**c**) and ligand-bound (**d**) CK-BB at physiological and acidic pH. The MD simulations reproduce the ligand- and pH-dependent opening observed in the DEER-guided structural ensembles.

**Figure 6.**
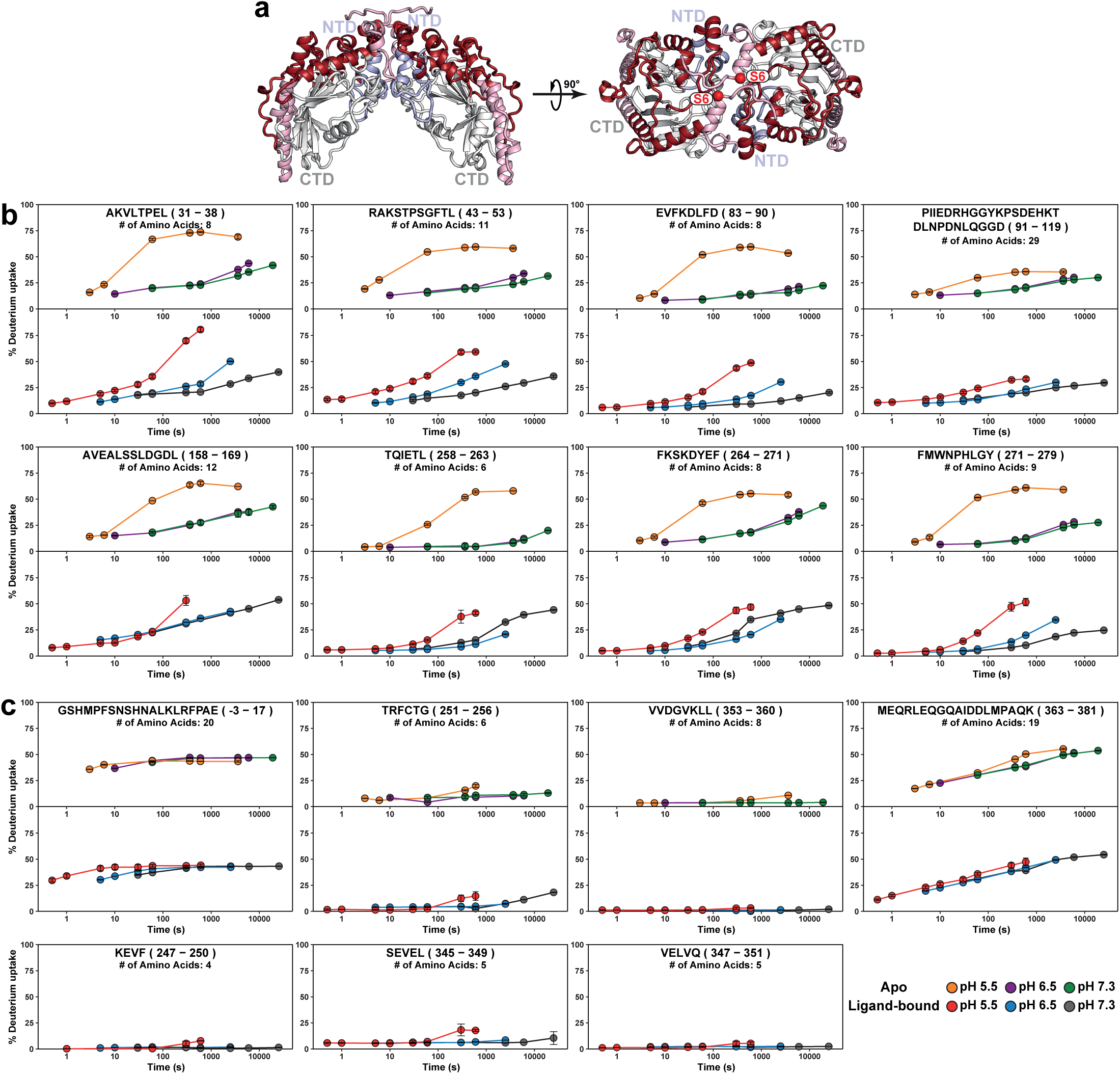
Acidic pH and ligand binding selectively enhance dynamics at the top of the CK-BB convex surface. (**a**) CK-BB dimer structure with convex-surface peptides mapped onto the NTD and CTD. Peptides showing increased deuterium uptake under acidic conditions are shown in dark red, whereas pH-invariant or minimally exchanging peptides are shown in light red; the Ser6 phosphorylation site is shown as a red sphere. (**b**) HDX kinetic plots for peptides near the top of the convex surface, excluding the N-terminal region, show pronounced pH-dependent increases in backbone exchange. In apo CK-BB, exchange is largely similar at pH 7.3 and 6.5 but increases markedly at pH 5.5, consistent with a threshold-like acid-induced transition. Ligand binding sensitizes this region to pH, producing modest but graded increases in HDX across all three pH conditions, consistent with allosteric, substrate-dependent priming of local backbone dynamics and/or solvent exposure. (**c**) In contrast, peptides along the lateral sides of the convex surface show minimal pH- or ligand-dependent changes, indicating comparatively stable or protected regions. Deuterium uptake is plotted as percentage uptake over exchange time for apo and ligand-bound CK-BB at pH 7.3, 6.5 and 5.5.

**Figure 7.**
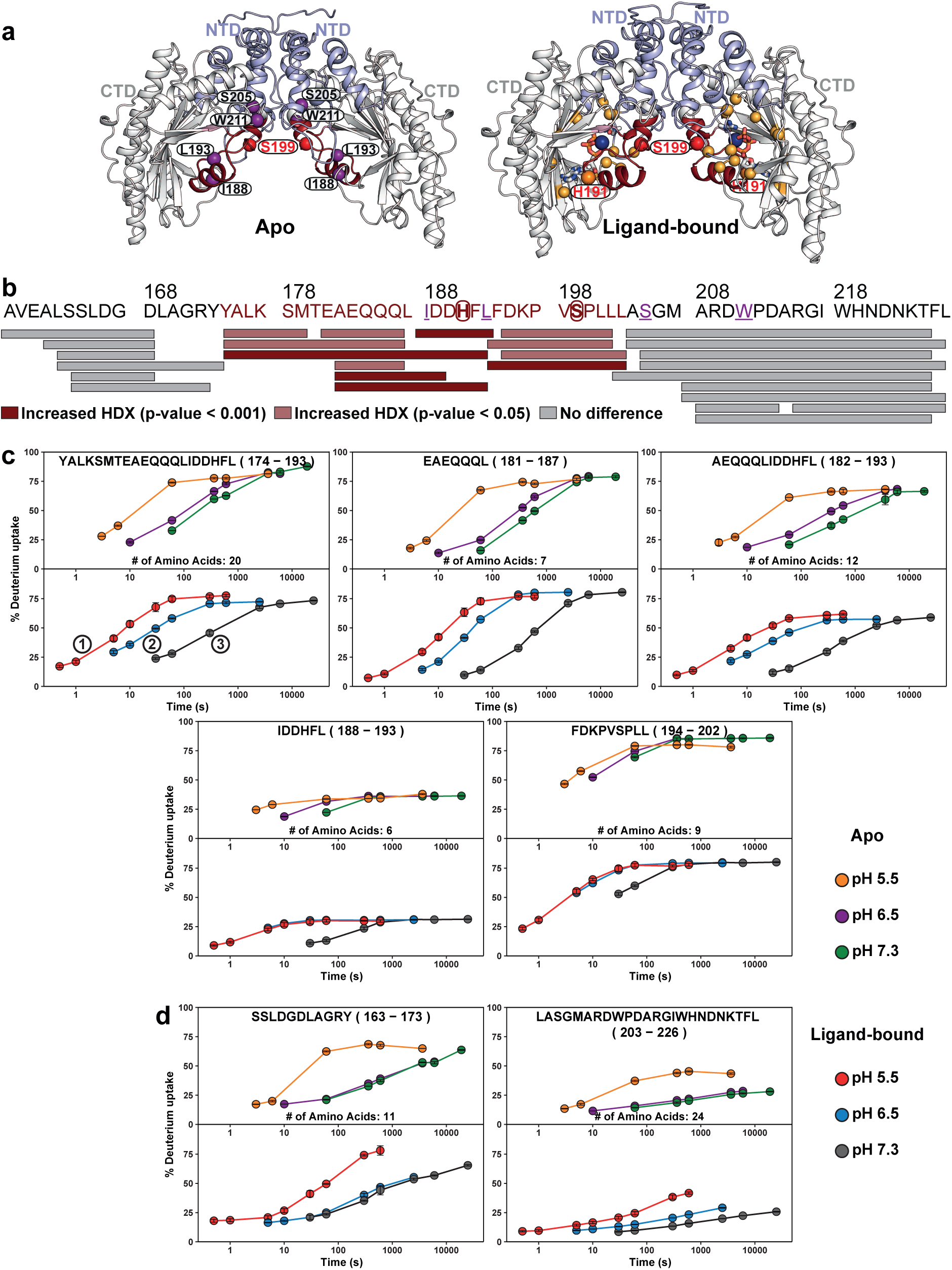
Ligand-sensitized, pH-dependent dynamics near His191 and Ser199 on the CK-BB concave surface. (**a**) Apo and ligand-bound CK-BB dimer models highlighting the concave-surface region spanning the nucleotide ribose-interacting residue His191, the putative Ser199 phosphorylation site, and associated DEER reporter positions (purple spheres, Fig. 4). Catalytic and substrate-binding residues are shown as orange spheres. (**b**) Expanded peptide coverage map for residues 158–226, highlighting peptides with significant pH-dependent increases in deuterium uptake. (**c**) HDX uptake plots for peptides spanning the His191/Ser199 region. In apo CK-BB, exchange at pH 6.5 remains closer to pH 7.3, whereas pH 5.5 markedly increases exchange, consistent with a threshold-like acid-induced enhancement of local backbone dynamics and/or solvent exposure. Ligand binding sensitizes this region to pH, producing graded increases in HDX across all three pH conditions, consistent with substrate-dependent priming of local dynamics. (**d**) HDX uptake plots for peptides flanking the His191/Ser199 region. Unlike the central region, these flanking peptides show primarily threshold-like acid-induced increases in exchange under both apo and ligand-bound conditions. Global significance threshold, 0.32 Da; *p* < 0.001.

These DEER-guided conformational trends were independently supported by molecular dynamics simulations. Across triplicate 1-µs simulations, dimer-opening angle distributions shifted toward larger opening angles under acidic conditions and upon ligand binding, with the most open conformations enriched in the ligand-bound acidic state (Figs. 5c and 5d). The convergence of DEER-guided ensembles, AF3-refined models and MD-derived opening angles indicates that CK-BB samples a continuum of pH- and ligand-dependent open states rather than two rigid endpoints, with protonation and ligand binding progressively biasing the landscape toward dimer expansion and hinge-like opening. This progressive opening parallels the membrane-association measurements, with substrate occupancy promoting CK-BB binding at neutral pH and acidification eliciting a stronger response. Together, these findings support a mechanism in which dimer opening shifts CK-BB toward membrane-competent conformations that increasingly favor membrane engagement.

### HDX-MS resolves region-specific acid-enhanced CK-BB dynamics

HDX-MS provides a complementary, label-free approach for mapping CK-BB dynamics because backbone amide exchange reports local hydrogen bonding, conformational dynamics, and solvent accessibility at peptide resolution^74–77^. Because intrinsic amide exchange is strongly pH-dependent, labeling times were adjusted across pH conditions before interpreting uptake differences as conformational or accessibility changes^58,59^. This makes HDX-MS particularly well suited to connect acid-induced structural remodeling with the emergence of membrane-competent CK-BB conformations^57–61^.

Native MS confirmed that CK-BB remains predominantly dimeric under the conditions used for structural analysis. The dominant native species was observed at 85,851 ± 1 Da, consistent with stable CK-BB dimer formation, whereas the smaller monomeric signal likely reflects partial gas-phase dissociation during ionization (Supplementary Figs. 8a and 8b). A minor +64 Da dimeric species was also detected, consistent with an unidentified post-translational or chemical modification. Denaturing MS yielded a monomeric mass of 42,925 Da, in close agreement with the calculated CK-BB mass of 42,925.5 Da, with minor higher-mass species at +98, +198 and +315 Da (Supplementary Figs. 8c and 8d). The absence of broad HDX protection expected for aggregation or precipitation further supports acidification-induced CK-BB redistribution rather than nonspecific aggregation.

**Figure 8.**
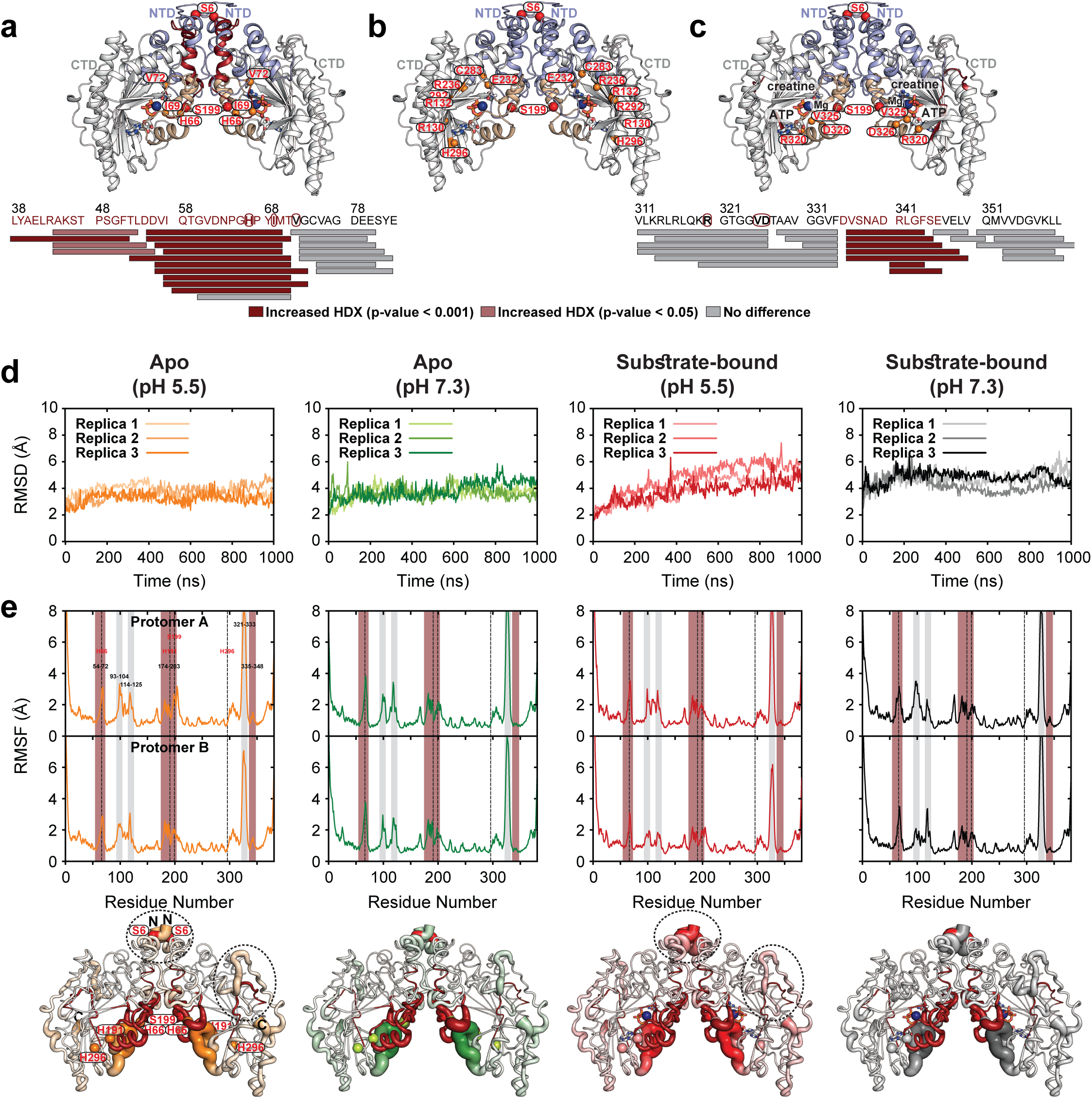
Molecular dynamics simulations link acid- and ligand-sensitive HDX regions to dynamic CK-BB structural elements. (**a**) CK-BB model highlighting the NTD interface and adjacent conserved flexible loop (residues 60–70), including His66 and Ile69, which show increased deuterium uptake under acidic conditions (Supplementary Fig. 11). Functional residues in this region are shown as orange spheres. (**b**) Additional catalytic and substrate-binding residues on the concave surface. (**c**) CK-BB model highlighting a C-terminal beta-sheet core region that shows increased deuterium uptake in the ligand-bound state under acidic conditions. (**d**) Cα RMSD traces from three independent 1-µs MD simulations of apo and ligand-bound CK-BB at pH 7.3 and pH 5.5, showing overall structural stability with greater conformational variability in the ligand-bound acidic state. (**e**) Per-residue root-mean-square fluctuation (RMSF) profiles for the same simulations. Shaded regions mark peptides with significant pH-dependent increases in deuterium uptake (red) and additional flexible segments (grey). Functional histidines and the Ser199 phosphorylation site are indicated by dashed lines. Bottom panels show RMSF values mapped onto CK-BB structures, with ribbon thickness proportional to local flexibility and HDX-sensitive regions highlighted in dark red. Together, the simulations support the HDX-MS data by showing that acid- and ligand-sensitive exchange regions either coincide with locally flexible segments or lie adjacent to dynamic elements that may modulate solvent exposure.

We used HDX-MS to localize regions of CK-BB that undergo pH-dependent changes in backbone exchange. Because intrinsic exchange slows at lower pH, labeling times were adjusted to compare equivalent intrinsic-exchange windows across conditions (Supplementary Table 1). Pepsin digestion yielded 170 identified peptides, 100% sequence coverage, an average peptide length of 13.4 residues and a redundancy of 6.4, supporting robust peptide-level analysis (Supplementary Fig. 9). Volcano plots and peptide coverage maps showed that decreasing pH progressively increased both the magnitude and statistical significance of deuterium uptake in apo CK-BB relative to pH 7.3 (Supplementary Figs. 10a–d). This response was modest and localized at pH 6.5, broader at pH 6.0 and widespread at pH 5.5. Ligand-bound CK-BB also showed acid-dependent increases in exchange, although the affected regions differed modestly from those in the apo state (Supplementary Figs. 10e and 10f).

These increases in deuterium uptake support condition-dependent changes in local backbone dynamics and/or solvent exposure. Acidification broadly enhanced CK-BB backbone exchange while preserving strong regional specificity. The top of the convex dimer surface showed pronounced pH-dependent increases in exchange, whereas peptides along the lateral convex surfaces remained comparatively protected (Fig. 6). In apo CK-BB, exchange across the top convex surface changed modestly at pH 6.5 but increased markedly at pH 5.5, consistent with a threshold-like acid-induced transition. Ligand binding sensitized this region to pH, producing modest but graded increases in exchange across pH 7.3, 6.5 and 5.5. This HDX-MS pattern closely parallels the DEER measurements, which show that convex-surface reporters remain comparatively restrained with limited pH- or ligand-dependent redistribution except under stronger acidification (Fig. 3).

### Ligand binding sensitizes the His191/Ser199 concave-surface module to pH

HDX-MS identified a second major pH-responsive region on the concave surface of CK-BB, centered on His191, which interacts with nucleotide ribose, and Ser199, a consensus ERK1/2 mitogen-activated protein kinase (MAPK) and putative AMP-activated protein kinase (AMPK) phosphorylation site (Fig. 7)^73,78^. His191 lies within the nucleotide-binding environment, whereas Ser199 is positioned adjacent to a short helix connected to the C-terminal β-sheet core (Fig. 7a). In apo CK-BB, peptides spanning this region showed only modest exchange differences between pH 7.3 and 6.5 but markedly increased deuterium uptake at pH 5.5, consistent with a threshold-like acid-induced increase in local backbone dynamics and/or solvent exposure.

Ligand binding reshapes this response. Rather than uniformly protecting the His191/Ser199 region, substrate binding sensitized it to pH, producing graded increases in exchange across pH 7.3, 6.5 and 5.5 (Fig. 7c). The strongest responses were observed in peptides spanning residues 174–193 and 181–193, whereas shorter peptides centered on residues 188–193 showed more limited changes, suggesting local protection within the His191-containing segment (Supplementary Fig. 14). Flanking peptides, including residues 158–173 and 203–226, also displayed acid-enhanced exchange (Fig. 7d and Supplementary Fig. 14), indicating that pH-dependent backbone dynamics extend beyond the central His191/Ser199 site across the surrounding concave surface.

Together with the DEER data (Fig. 4), which show pH- and ligand-dependent redistribution of conformational intermediates across this concave face, these HDX-MS results identify the His191/Ser199 region as a ligand-sensitized regulatory module in which protonation and substrate occupancy are coupled to inter-protomer ensemble remodeling.

### Molecular dynamics connect acid-sensitive HDX regions to dynamic structural elements

To relate HDX-sensitive regions to intrinsic flexibility or acidification- and ligand-dependent structural remodeling, we performed triplicate 1-µs molecular dynamics simulations of apo and ligand-bound CK-BB at pH 7.3 and pH 5.5. Across all four states, Cα root-mean-square deviation (RMSD) traces showed overall structural stability, with the greatest conformational variability observed in the ligand-bound acidic state (Fig. 8d). Per-residue root-mean-square fluctuation (RMSF) profiles further showed that many acid- and ligand-sensitive HDX regions either coincide with locally flexible segments or lie adjacent to dynamic elements that could modulate solvent exposure (Fig. 8e). For example, residues 335–348, which lie within the largely buried CTD β-sheet core, are adjacent to a region with increased dynamics under acidic conditions (circled, Fig. 8e), potentially enhancing their solvent accessibility. The NTD– NTD dimer interface becomes more dynamic in the ligand-bound acidic state than in the apo state, potentially promoting sampling of more open ligand-bound conformations.

Three regions are particularly informative. First, the conserved flexible loop near the NTD interface, containing His66, Ile69 and Val72, showed acid-enhanced deuterium uptake and mobility in MD simulations (Figs. 8a and 8e and Supplementary Figs. 11–13). Peptides spanning residues 52–73 and 72–90 showed a threshold-like acid-induced increase in exchange in apo CK-BB, whereas ligand-bound CK-BB exhibited more graded pH-dependent exchange across overlapping peptides. This ligand-sensitized behavior extended to regions containing Arg130, Arg132, Glu232, Arg236, Cys283, Arg292 and His296 (Supplementary Fig. 11), indicating that substrate binding tunes the pH responsiveness of catalytic and substrate-binding elements.

Second, the concave catalytic surface, including His191/Ser199 and neighboring Glu232, Cys283, Arg292 and His296, contained pH-responsive segments (Figs. 7 and 8b and Supplementary Figs. 14–16). Residues 207–226 and 251–281 showed acid-enhanced exchange, whereas 226–250, including Glu232/Arg236, remained protected. Thus, acidification selectively increases exchange in discrete elements adjacent to catalytic and ligand-binding residues rather than globally destabilizing the active site.

Third, the C-terminal β-sheet core containing Arg320, Val325 and Asp326 showed heterogeneous pH sensitivity (Fig. 8c and Supplementary Fig. 17). Residues 318–334 exhibited high baseline exchange with limited condition dependence, 335–348 showed modest acid sensitivity, and 363–381 were pH- and ligand-invariant. Together, the MD and HDX-MS analyses show that acid- and ligand-sensitive exchange is localized to discrete dynamic modules rather than distributed uniformly across CK-BB.

### Ligand binding buffers global acid-induced deprotection while preserving local pH sensitivity

To compare apo and ligand-bound states directly, we mapped deuterium uptake differences at 60 s onto CK-BB structures and peptide-level bar plots (Fig. 9). In apo CK-BB, acidic pH broadly increased exchange across the NTDs, top convex surface and His191/Ser199-containing concave region (Figs. 9a, 9c, and 9e). Ligand binding substantially attenuated this global acid-induced deprotection, particularly across the otherwise dynamic convex and NTD regions (Figs. 9b, 9e, and 9h). For this direct comparison, the analysis was restricted to the 120 peptides that were detected and matched across both apo and ligand-bound datasets.

**Figure 9.**
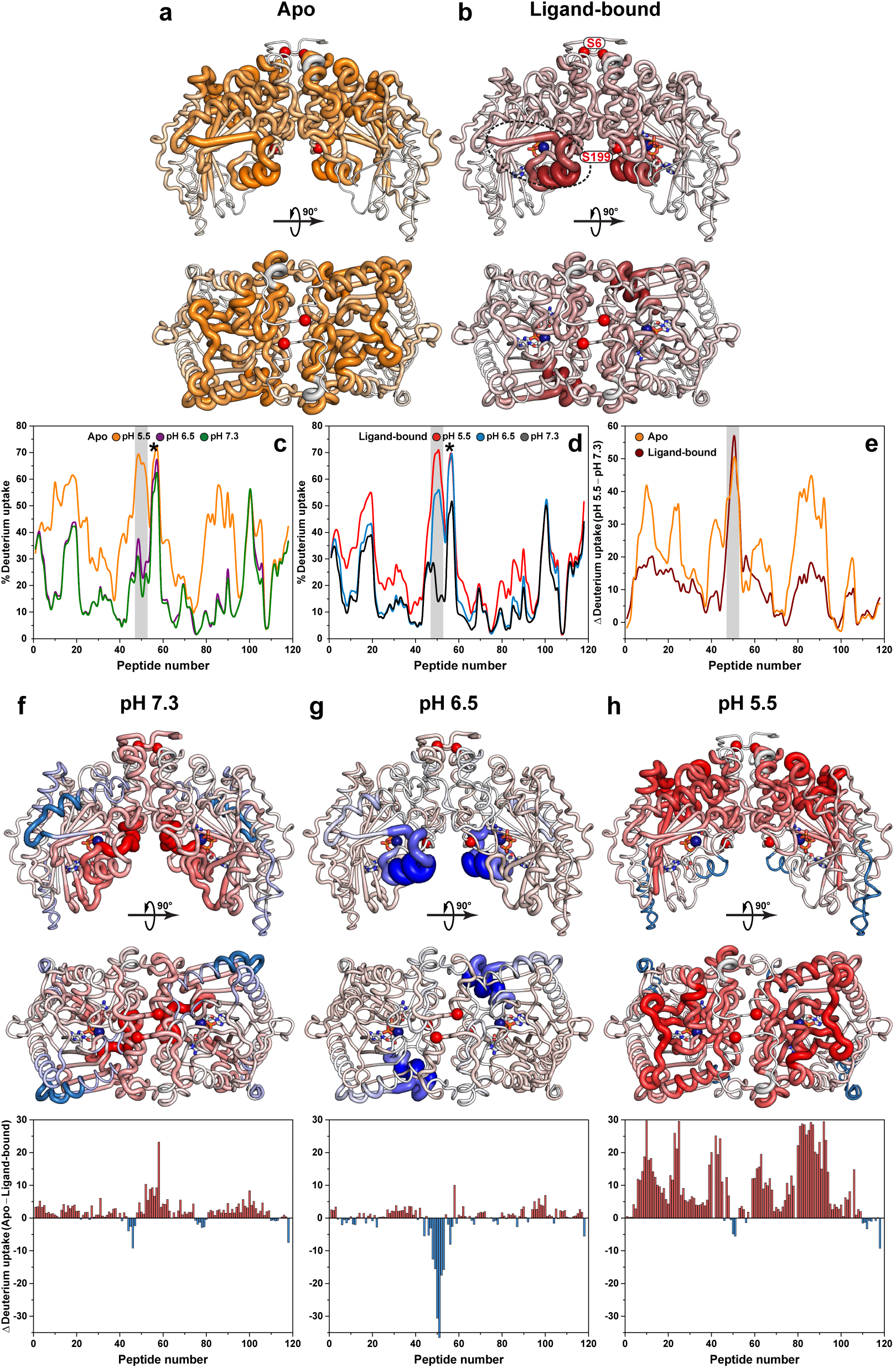
Ligand binding buffers global acid-induced CK-BB dynamics while sensitizing the His191/Ser199 region. (**a**,**b**) Differences in deuterium uptake at 60 s between pH 5.5 and pH 7.3 for apo (**a**) and ligand-bound (**b**) CK-BB, mapped onto the CK-BB dimer with ribbon thickness proportional to increased uptake under acidic conditions. In apo CK-BB, acidic pH broadly increases deuterium uptake across the NTDs, convex surface, and the His191/Ser199-containing concave region, whereas ligand binding substantially attenuates this global acid-induced deprotection while preserving a localized response near His191/Ser199. (**c**,**d**) Peptide-level uptake profiles across pH for apo (**c**) and ligand-bound (**d**) CK-BB. The shaded region marks peptides surrounding the short helix adjacent to the C-terminal beta-sheet core and preceding the Ser199 phosphorylation site (circled in **b**). In apo CK-BB, exchange at pH 6.5 remains closer to pH 7.3, whereas pH 5.5 produces a threshold-like increase in uptake. Ligand binding sensitizes this region to pH, producing graded increases in exchange across all three pH conditions. (**e**) Direct comparison of acid-induced uptake changes in apo and ligand-bound CK-BB highlights the global protective effect of ligand binding and the retained pH sensitivity of the His191/Ser199 region. (**f**–**h**) Differences in deuterium uptake at 60 s between apo and ligand-bound CK-BB at pH 7.3, 6.5 and 5.5, mapped onto the CK-BB dimer and shown as peptide-level bar plots. Positive values indicate greater uptake in apo CK-BB, consistent with ligand-dependent protection; negative values indicate greater uptake in the ligand-bound state. At pH 5.5, ligand binding strongly protects otherwise dynamic convex and NTD regions.

However, ligand binding did not uniformly suppress pH sensitivity. The His191/Ser199 region retained a localized acid response (Fig. 9b), and the short helix preceding Ser199 became more sensitive to graded pH changes in the ligand-bound state (shaded region, Figs. 9c–9e). Positive differences at pH 5.5 indicated that ligand binding protects large portions of CK-BB from acid-induced exchange (Fig. 9h), whereas localized negative or attenuated differences identified regions that remain dynamic or become selectively sensitized upon ligand binding (Fig. 9f-h). Thus, substrate binding stabilizes the global dimer while priming a regulatory concave-surface region for pH-dependent remodeling.

## Discussion

This study identifies progressive dimer opening as a novel regulatory mechanism that couples protonation and substrate binding to membrane engagement by human CK-BB. Across cellular imaging, tethered-vesicle binding, DEER spectroscopy, HDX-MS, DEER-guided modeling and MD simulations, we show that CK-BB occupies an acid- and substrate-regulated conformational landscape in which dimer opening drives region-specific remodeling of the convex and concave surfaces. Acidification drives curvature-sensitive association with plasma membrane and synaptic vesicle mimetics, redistributes CK-BB to vesicular structures and membrane ruffles, and increases backbone exchange across selected regions, whereas substrate binding independently promotes membrane association at neutral pH. This opening is coupled to lateral dimer expansion (PC1, Supplementary Video 1), with ligand-bound CK-BB at pH 5.5 occupying the most laterally expanded and open region of the conformational landscape. Substrate binding buffers broad acid-induced deprotection while preserving and sharpening local pH sensitivity near the His191/Ser199 catalytic-regulatory interface. Together, these findings define dimer opening as a switch that generates membrane-competent CK-BB states and couples local ATP regeneration to curved, actin-remodeling membrane domains.

A central insight is that CK-BB dynamics are spatially asymmetric. The convex face of the dimer, including the NTD interface and C-terminal β-sheet core, remains comparatively restrained by DEER under most conditions except strong acidification; consistently, the top of this surface becomes HDX-sensitive upon acidification. Because HDX labeling times were pH-adjusted, greater uptake at acidic pH reflects region-specific structural remodeling. Substrate binding further converts the threshold-like apo response into a more graded pH-sensitive transition.

The concave His191/Ser199 region behaves differently. DEER measurements reveal pH- and ligand-dependent redistribution among multiple conformational intermediates, whereas HDX-MS shows ligand-sensitized increases in local backbone exchange near His191 and Ser199. His191 lies within the nucleotide-binding environment, while Ser199 occupies a regulatory surface adjacent to a short helix connected to the CTD core. In the region preceding the phosphorylation site, highlighted by the grey shading and asterisk in Figs. 9c–9e, apo CK-BB displays a threshold-like acid response, whereas ligand-bound CK-BB shows a more graded pH-dependent increase in exchange. Thus, DEER captures large-scale inter-protomer remodeling of the concave surface, while HDX-MS resolves accompanying local changes in backbone dynamics and/or solvent exposure. Their convergence identifies the His191/Ser199 region as a ligand-sensitized, acid-responsive regulatory module through which active-site occupancy is communicated to dimer-scale opening. Notably, in the ligand-bound state, the region surrounding the phosphorylation site, which is coupled to the NTD–NTD interface, becomes more dynamic or exposed even under mildly acidic conditions (pH 6.5). These dynamics could facilitate Ser199 phosphorylation by ERK1/2 MAPK or AMPK-linked stress-responsive pathways. Consistent with this possibility, AMPK responds to metabolic stress and phosphagen balance, such that reduced PCr, increased creatine and mild acidification can promote its activation^73^.

The peptide-resolved HDX and MD data further refine this mechanism. The conserved flexible loop containing His66, Ile69 and Val72 is strongly acid-sensitive, consistent with prior structural and mutational work assigning this loop to active-site closure and catalytic organization^42,44,45^. Substrate binding extends this pH sensitivity to additional functional regions, including peptides containing Arg130, Arg132, Glu232, Arg236, Cys283, Arg292 and His296, indicating that ligand occupancy broadly tunes the pH responsiveness of catalytic and substrate-binding elements. Several peptides spanning the Glu232/Arg236 region and portions of the CTD core remain comparatively protected, showing that acidification does not globally destabilize CK-BB. Instead, acidic pH selectively enhances exchange in discrete dynamic modules while preserving protected catalytic and core regions. This organization may allow CK-BB to maintain catalytic competence while increasing the conformational flexibility required for localization or membrane-proximal function.

These findings provide a structural rationale for the long-standing connection between CK-BB and actin-rich membrane compartments. Previous cell-biological studies showed that CK-B accumulates in membrane ruffles and phagocytic cups, where local CK activity supports ATP-dependent actin remodeling^9,10,26^. Our imaging data extend this framework by showing that acidification alone redistributes endogenous and recombinant CK-BB from a diffuse cytosolic pool to punctate vesicular structures and membrane ruffles. Tethered-vesicle measurements demonstrate that acidification directly promotes CK-BB membrane association, while substrate availability independently enhances binding at neutral pH. These membrane responses closely parallel conformational remodeling: substrate binding promotes dimer opening at neutral pH, whereas acidification elicits stronger membrane binding and shifts CK-BB further toward open states. Together, these findings support a model in which progressive dimer opening remodels the dynamic convex and concave surfaces, exposing or stabilizing membrane-competent conformations that enable a soluble enzyme lacking a canonical membrane anchor to accumulate at curved membrane surfaces.

Membrane curvature provides an additional layer of regulation. CK-BB binding density increased markedly as vesicle diameter decreased under acidic conditions, with even stronger curvature sensitivity on synaptic vesicle-mimetic membranes; substrate-dependent binding at neutral pH showed a similar preference for highly curved surfaces. The crescent-shaped CK-BB dimer presents distinct convex and concave faces, raising the possibility that pH- and substrate-dependent remodeling of these surfaces tunes their capacity to engage vesicular or actin-remodeling membranes. Although the precise membrane-binding interface and lipid determinants remain unresolved, our measurements directly establish curvature-sensitive CK- BB association and identify acid- and ligand-responsive surfaces that could contribute to this behavior. FRAP further showed that most membrane-bound CK-BB remained associated over several minutes, consistent with a persistent, slowly exchanging population. These findings identify CK-BB as a previously unrecognized curvature-sensitive metabolic enzyme and extend membrane-curvature sensing to the creatine kinase family. Together with longstanding evidence that other creatine kinase isoforms engage membranes, they raise the possibility that curvature-dependent membrane recognition may represent a broader, isoform-specific feature of creatine kinase biology.

Integrating these observations, we propose a model in which acidic pH and substrate binding remodel CK-BB to support membrane-proximal function (Fig. 10). Progressive dimer opening acts as a regulatory switch that increases dynamics across the top convex surface and NTD–NTD interface, sensitizes the His191/Ser199-containing concave region, and generates membrane-competent CK-BB conformations. These changes may help the dimer adapt to curved vesicular and cortical membrane surfaces formed during ruffling, endocytic recycling or vesicle-associated trafficking. Locally recruited CK-BB could then regenerate ATP at sites of membrane remodeling, supporting actin polymerization, membrane protrusion, and vesicle-associated dynamics. Thus, CK-BB emerges as an acid- and substrate-regulated metabolic module whose conformational state couples local ATP buffering to dynamic membrane compartments.

**Figure 10.**
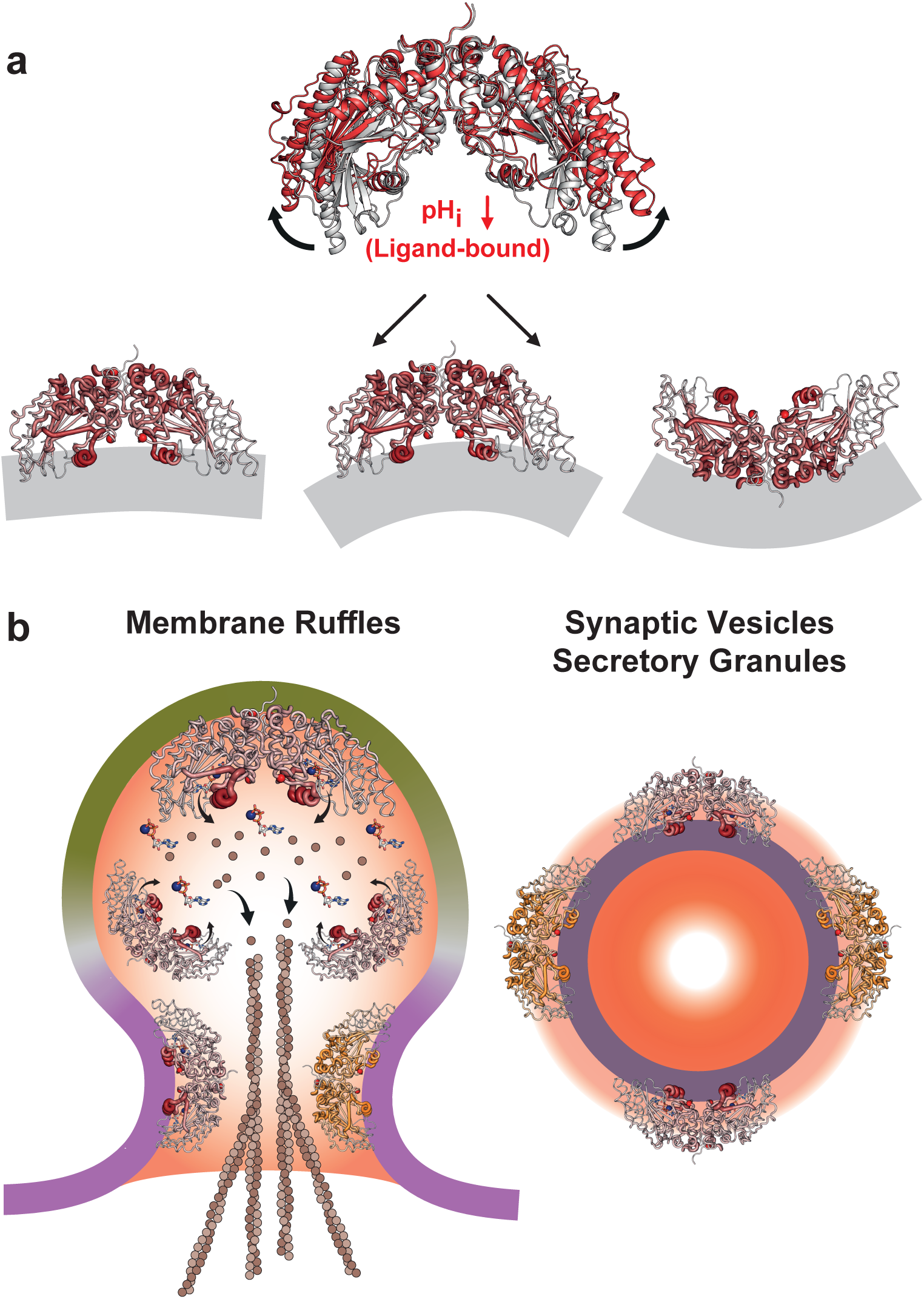
Model for pH- and ligand-regulated CK-BB curvature sensing and membrane coupling. Integrated model linking CK-BB conformational remodeling to curvature-sensitive membrane association and recruitment to vesicular and actin-remodeling membranes. (**a**) DEER, HDX-MS and MD support a mechanism in which acidification, particularly in the ligand-bound state, promotes progressive dimer opening and increases dynamics across the convex surface and His191/Ser199-containing concave region. Tethered-vesicle measurements show that acidification and substrate binding independently promote CK-BB association with highly curved plasma membrane and synaptic vesicle mimetics, supporting a model in which dimer opening generates membrane-competent conformations that favor curvature sensing and membrane engagement. (**b**) In cells, acidification redistributes CK-BB from a largely cytosolic pool to punctate vesicular s1tructures and membrane ruffles, where locally recruited CK-BB may regenerate ATP to support actin remodeling, membrane protrusion and vesicle-associated dynamics. Together, these findings link CK-BB conformational regulation, membrane-curvature sensing and localized ATP buffering at dynamic membrane compartments.

This mechanism may be relevant to cancer and stress physiology. Hypoxic and metabolically stressed tumor cells can release CK-BB, and extracellular CK-BB has been linked to phosphocreatine generation, anoikis resistance, hypoxic survival and metastatic progression^11,13,16^. Recent work connects creatine-phosphagen metabolism to pancreatic cancer invasion, SLC6A8-dependent metabolic vulnerabilities, glioblastoma growth in hypoxic niches, CK-dependent tumor-cell migration and invasion, and ferroptosis suppression through a moonlighting GPX4-phosphorylation function^18–23^. Because acidic pH is common in hypoxic and metabolically active microenvironments, and local acidification accompanies endosomal, vesicular, and secretory pathways, a pH- and substrate-responsive CK-BB dimer may provide a molecular route by which metabolic stress promotes CK-BB relocalization, membrane association, and extracellular availability. The ability of substrate binding to buffer global deprotection while preserving localized dynamics may help maintain enzymatic competence during membrane engagement. It may also be relevant to CK-BB dysfunction in neurodegeneration, where altered pH homeostasis, oxidative modification and aberrant membrane partitioning could perturb the balance between catalytic activity and membrane-proximal localization^30–32^.

The strength of the current model lies in the convergence of orthogonal approaches spanning quantitative membrane binding and curvature sensing, DEER, HDX-MS, molecular dynamics, DEER-guided modeling and cellular redistribution. Future studies should define the molecular architecture of the CK-BB–membrane interface, resolve contributions of specific lipid species, and determine how membrane engagement intersects with catalytic regulation and phosphorylation at Ser6 and Ser199. Together, these results establish dimer opening as a novel regulatory mechanism that tunes CK-BB for local ATP buffering at dynamic membranes. This structural-dynamics framework connects catalytic regulation, vesicular redistribution and actin-remodeling microdomains, establishes CK-BB as a previously unrecognized curvature-sensitive metabolic enzyme, and provides a foundation for understanding its contributions to cancer metabolism, extracellular phosphocreatine production and stress-associated membrane remodeling.

## Methods

No statistical methods were used to predetermine sample size. Experiments were not randomized. Investigators were not blinded to allocation during experiments or outcome assessment.

### Cell culture, acidification and immunofluorescence

HCT116, HCT116/FLAG-CK-BB, HEK293 and HEK293/FLAG-CK-BB cells were cultured overnight on glass-bottom chambers coated with 2 µg cm^-2^ recombinant human laminin-521 (Gibco, A29248). Cells were treated for 1 h in Krebs–Ringer Bicarbonate HEPES buffer (KRBH) containing 129 mM NaCl, 4.8 mM KCl, 1.2 mM KH_2_PO_4_, 1.2 mM MgSO_4_, 2.5 mM CaCl_2_, 20 mM HEPES and NaHCO_3_ as indicated. Acute extracellular acidification was used as a stress-like perturbation to test CK-BB redistribution. Target pH values were set by titration before application to cells. To maintain pH during the 1-h treatment in a 5% CO_2_ incubator, neutral pH 7.2–7.4 medium was prepared in KRBH containing 20 mM NaHCO_3_, whereas acidic pH 5.5 medium was prepared in KRBH without NaHCO_3_, consistent with published guidelines^79^. Cells were fixed with 4% paraformaldehyde for 30 min at room temperature, permeabilized with 0.1% Triton X-100 in PBST (PBS containing 0.05% Tween 20) for 10 min at room temperature, and blocked for 1 h in PBST containing 1% BSA and 5% goat serum. Cells were incubated overnight at 4 °C with anti-CK-BB antibody (Proteintech, 15137-1-AP) or anti-FLAG antibody (Sigma, F3165) diluted in PBST containing 1% BSA, followed by 1 h incubation with goat anti-rabbit Alexa Fluor 488 or goat anti-mouse Alexa Fluor 594 secondary antibodies. Images were collected using a Zeiss LSM880 fluorescence microscope.

### Site-directed mutagenesis

Human CK-BB gene was cloned into pET28b, encoding an N-terminal 6-His tag under an inducible T7 promoter. The five cysteine residues in CK-BB were mutated (C74A, C141T, C146S, C254S and C283D) by site-directed mutagenesis using complementary oligonucleotide primers, yielding the CL protein. This construct served as the template for single-cysteine DEER and fluorescence variants. Substitutions were generated by single-step PCR using a mutagenic primer. Mutants were verified by sequencing with T7 forward and reverse primers. Mutants were identified by the native residue and primary sequence position followed by the substituted residue.

### Expression, purification, and labeling of CK-BB

*Escherichia coli* BL21(DE3) cells were transformed with pET28b encoding WT or mutant CK-BB. A single colony was grown overnight (∼15 h) at 37 °C in Luria–Bertani (LB) medium (Fisher Bioreagents) containing 0.05 mg/mL kanamycin (Gold Biotechnology) and used to inoculate 3–6 L LB at 1:50. Cultures were shaken at 37 °C to OD600 ∼0.5–0.7 and induced with 0.25 mM IPTG (Gold Biotechnology). Cultures were incubated for 4 h at 30 °C and harvested by centrifugation. Cell pellets were resuspended in lysis buffer (20 mM HEPES, pH 7.5, 300 mM NaCl, and 5% [vol/vol] glycerol) at 25 mL/L culture, supplemented with 2 mM DTT and 1 mM PMSF, and sonicated for 8 min total using 15-s on/15-s off pulses at 30% amplitude. Cell debris was removed by centrifugation at 30,000 r.p.m. (105,000 × *g*) for 30 min at 4 °C in a Type 45 Ti rotor (Beckman Coulter). The supernatant was incubated with 2.0 mL (bed volume) His60 Ni Superflow resin (TakaraBio) for 2 h at 4 °C, washed with 10 column volumes of buffer containing 30 mM imidazole and eluted with buffer containing 300 mM imidazole. The eluate was desalted into PBS containing 5% glycerol using a Cytiva HiTrap desalting column. The His tag was removed by thrombin protease (MP Biochemicals) overnight at 4 °C at a 1:1000 thrombin:CK-BB mass ratio (ε_CK-BB-CL_ = 35,410 M^−1^⋅cm^−1^, M.W. ∼ 44,753 Da). For DEER spectroscopy and fluorescence microscopy, single-cysteine mutants were labeled, respectively, with a 10-fold molar excess per cysteine of 1-oxyl-2,2,5,5-tetramethylpyrroline-3-methyl methanethiosulfonate (Enzo Life Sciences) or AZDye 488 Maleimide (Vector Laboratories). Thrombin was removed using a Cytiva HiTrap Heparin HP column (20 mM Tris pH 7.5, 5% glycerol), and cleaved CK-BB was separated from uncleaved protein and free His tag using 2 mL Ni resin on a gravity flow column while collecting cleaved CK-BB in the flow-through. Final purification was performed by size-exclusion chromatography on a Superdex 200 Increase 10/300 GL column (Cytiva) equilibrated in 50 mM Tris/MES, pH 7.3, 100 mM NaCl buffer, with or without 10% (vol/vol) glycerol buffer for DEER spectroscopy and MS analyses, respectively. Peak fractions were pooled and concentrated using a 30-kDa MWCO Amicon Ultra concentrator (Millipore, Sigma-Aldrich). The final concentration was determined by A_280_ measurement (ε = 35,410 M^−1^⋅cm^−1^, M.W. ∼ 42,871 Da) for use in subsequent studies. WT CK-BB was analyzed by native and denaturing MS to confirm its molecular mass.

### Creatine kinase activity assay

Creatine kinase activity was measured using the creatine kinase activity kit (MAK116, Sigma-Aldrich). Recombinant CK-BB was diluted to 0.1 mg/mL in 50 mM Tris, pH 7.5, and 1 µg protein was used per reaction. Reactions were initiated by adding reconstituted kit reagent containing assay buffer, enzyme mix, and substrate solution. For pH-dependent assays, the reagent was prepared in 50 mM Tris/MES and 150 mM NaCl buffers adjusted to the indicated final pH values. Absorbance at 340 nm was monitored every minute for 40 min at 37 °C on a SpectraMax i3 plate reader. Activity was determined from the linear change in NADH-coupled absorbance according to the manufacturer’s equation. Experiments were performed in ≥3 replicates, and mean ± s.d. was calculated.

### Circular dichroism spectroscopy

Far-UV circular dichroism spectra were recorded for WT and CL CK-BB, and for spin-labeled mutants. Proteins were diluted to 0.01 mg/mL in 10 mM sodium phosphate, pH 7.3. Spectra were collected from 195 to 260 nm at 22 °C using an Applied Photophysics Chirascan V100 in a 10-mm quartz cell. Data were collected at 1-nm intervals with 0.1 s per point and 10 averaged scans. Experiments for WT and CL CK-BB were performed in triplicate, and mean ± s.d. values were calculated.

### Vesicle preparation and tethering

1-palmitoyl-2-oleoyl-glycero-3-phosphocholine (POPC), 1-palmitoyl-2-oleoyl-glycero-3-phospho-L-serine (POPS), 1,2-dioleoyl-sn-glycero-3-phosphoethanolamine (DOPE), 1-stearoyl-2-arachidonoyl-sn-glycero-3-phosphoinositol (SAPI), and cholesterol (plant) were purchased from Avanti Research. Biotin-PEG-silane (5 kDa) and mPEG-silane (5 kDa) were purchased from Laysan Bio. Plasma membrane-mimetic vesicles contained 13.5% POPC, 34.5% DOPE, 22% POPS, 9% SAPI, 20% cholesterol, 0.5% DP-EG15-biotin (provided by D. Sasaki), and 0.5% ATTO 647N-DPPE (Leica Microsystems). Synaptic vesicle mimetics contained 24.5% POPC, 24.5% DOPE, 10% POPS, 40% cholesterol, 0.5% DP-EG15-biotin, and 0.5% ATTO 647N-DPPE. Lipid mixtures were dried under nitrogen, vacuum-desiccated for ≥2 h, rehydrated in 20 mM sodium phosphate and 100 mM NaCl, pH 7.3, subjected to five freeze–thaw cycles (1 min in liquid nitrogen; 5 min at 50 °C), and extruded through 50-nm polycarbonate membranes (Sterlitech). Plasma membrane mimetics were handled under nitrogen to minimize oxidation of polyunsaturated SAPI.

Slides were cleaned as described previously^71,72,80–82^ by sonication in 70% ethanol and sequential 10-min KOH/peroxide and HCl/peroxide treatments at 80 °C. Cleaned slides were passivated with 55 μL of PEG-silane solution containing biotin-PEG-silane and mPEG-silane in isopropanol, with glacial acetic acid added to catalyze silane coupling, and incubated for 1 h at 50 °C. Silicone gaskets (5-mm wells; Grace Bio-Labs) were applied to the passivated slides. Each well was incubated with 30 μL of 3.33 μM NeutrAvidin for 20 min, rinsed, and incubated with 1 μM vesicles for 10 min before removal of unbound vesicles. For acidic conditions, wells were exchanged into 20 mM sodium phosphate and 100 mM NaCl, pH 5.5. CK-BB, with or without substrates, was added at the indicated concentrations and incubated for 1 h before imaging (see Supplementary Methods).

### Confocal microscopy, FRAP, and image analysis

Images were acquired on a Leica Stellaris 5 laser-scanning confocal microscope equipped with a 63× oil-immersion objective (Leica HC PL APO, NA 1.4). AZDye 488-labeled CK-BB was excited at 488 nm and detected at 493–558 nm, whereas ATTO 647N-labeled vesicles were excited at 638 nm and detected at 643–749 nm. Images were collected at 70 × 70 nm pixels and a line-scanning frequency of 400 Hz. FRAP measurements used three pre-bleach images acquired over 2 s, five bleach frames at 100% laser power (∼4 s), and 50 recovery images at 5-s intervals.

Fluorescent puncta were identified and tracked using cmeAnalysis^72,80,81^. Vesicles detected in three consecutive frames and <250 nm in diameter were retained for analysis, and CK-BB binding was quantified from colocalized protein-channel intensity. Vesicle diameter was calibrated by matching the square root of the fluorescence-intensity distribution to the corresponding diameter distribution measured by dynamic light scattering. For FRAP analysis, vesicles were grouped into 40–60, 90–110, and 140–160 nm diameter bins for the nominal 50-, 100-, and 150-nm populations, respectively. Single-molecule AZDye 488 fluorescence was calibrated using CK-BB adsorbed to glass coverslips, with the peak of the resulting intensity distribution assigned to a single labeled CK-BB molecule and used to determine the number of membrane-bound proteins per vesicle. Calibration values were adjusted as necessary for differences in detector gain between imaging experiments (see Supplementary Methods).

### CW EPR and DEER spectroscopy

Room-temperature continuous-wave (CW) EPR spectra of spin-labeled CK-BB were collected on a Bruker EMX spectrometer at X-band (9.5 GHz) using 10-mW incident power and 1.6-G modulation amplitude to assess labeling quality. DEER spectroscopy was performed on an Elexsys E580 EPR spectrometer operating at Q-band frequency (33.9 GHz) with the dead-time free four-pulse sequence at 83 K^83^. Pulse lengths were 20 ns (π/2) and 40 ns (π) for the probe pulses and 40 ns for the pump pulse. The frequency separation was 63 MHz. To assess protonation effects, samples were titrated with HCl to pH 6.5 or 5.5 and confirmed using a pH microelectrode (Mettler Toledo InLab Ultra-Micro-ISM). The substrate-bound state was generated by addition of 10 mM MgATP, 0.8 mM ADP, 20 mM PCr and 10 mM Cr. Samples for DEER analysis were cryoprotected with 24% (vol/vol) glycerol and flash-frozen in liquid nitrogen. Primary DEER decays were globally analyzed across conditions for each spin pair using custom MATLAB (MathWorks) software as described^84^. Briefly, the software carries out global analysis of the DEER decays obtained under different conditions for the same spin-labeled pair. The distance distribution was assumed to consist of a sum of Gaussians, the number and population of which were determined based on a statistical criterion. Confidence bands were derived from fit-parameter uncertainties. Individual DEER-decay analysis yielded distributions consistent with the global analysis. Comparison of the experimental distance distributions with the CK-BB models and structures using a rotamer library approach was facilitated by the MMM software package^85^. Rotamer library calculations were conducted at 175 K.

### DEER-guided AlphaFold modeling

DEER-guided modeling of CK-BB was performed using DEERFold^53^, a fine-tuned AlphaFold2-based^55^ network that incorporates experimental DEER distance distributions as pairwise distograms within the Evoformer module, together with the multiple sequence alignment (MSA), to bias structure prediction toward spin-label-constrained conformations. The DEERFold architecture, training strategy, and input-encoding scheme were described previously^53^ and were used here without modification.

The CK-BB amino acid sequence was used as input, and its MSA was generated with the ColabFold MSA pipeline^86^. For DEERFold predictions, the MSA was subsampled to an effective sequence depth of *N*_eff_ = 10^87^. For each restraint condition described below, 50 independent models were generated using distinct random seeds, producing a conformational ensemble for that condition. Models were ranked by the Earth Mover’s Distance (EMD) between the predicted and experimental distance distributions, and the highest-ranked subset, hereafter referred to as the “top models,” was retained for downstream structural analysis.

For each spin pair, single-Gaussian approximation, defined by its mean distance and standard deviation, of the stabilized distance component observed under each condition was used as DEERFold restraints. Each single-Gaussian restraint set was supplied to DEERFold as an independent input, generating separate ensembles. Principal component analysis was performed on refined ensembles to identify dominant collective motions. PC1 reports lateral dimer expansion, whereas PC2 reports hinge-like opening and closure of the dimer.

### DEER-guided refinement of the unbiased AlphaFold3 models

Unbiased models of CK-BB in the apo and MgATP-bound states were generated using AlphaFold3 (AF3)^56^, as implemented in the AF3 server. Five independent runs were performed for each state, and the highest-pLDDT model was selected as the reference conformation. The

DEER distance constraints were used to refine the AF3-generated models using a previously published approach^54^. Refinement was carried out iteratively in MODELLER^88^. In silico spin labeling was performed with the MMM software package^85^ using a rotamer-library approach. In each iteration, rotamer ensembles were first calculated at 298 K for the spin-labeled sites, and the rotamer that best matched the mean N–O midpoint position of the full ensemble was attached to the template structure provided to MODELLER. Refinement was performed using the centers of the Gaussian components corresponding to each conformational state, together with secondary-structure restraints derived from AlphaFold models. All refined models achieved a GA341 score of 1.0. After import into MMM, rotamer ensembles were recalculated (Supplementary Figs. 6 and 7), and models were ranked by the root-mean-square deviation between the experimental distance constraints and the corresponding model-derived distances across all restraints.

### Hydrogen–deuterium exchange mass spectrometry

Bottom-up HDX-MS experiments were performed following established peptide-level HDX-MS principles^74–77^. Briefly, 5 µL of 20 µM CK-BB was diluted into 95 µL deuterated labeling buffer containing 25.6 mM Tris, 24.4 mM MES and 100 mM NaCl at the indicated pH values. Samples were labeled in triplicate at 20 °C for each pH-adjusted labeling time. Exchange was quenched by adding 100 µL quench buffer containing 100 mM glycine and 4 M urea, pH 2.4, precooled to 0 °C. Nondeuterated controls were prepared identically using H_2_O-based labeling buffer. For deuterated labeling buffers, pD was calculated from the apparent pH as pD = apparent pH + 0.4. For each labeling condition, the final quenched protein–buffer mixture was measured, and the quench composition was adjusted as needed to ensure equivalent final quench pH across all conditions. HDX-MS measurements were performed in triplicate at each labeling time and experimental condition (*n* = 3 technical replicates). Data are presented as the mean ± s.d., where applicable.

After quenching, 200 µL of each sample was injected using a fully automated Trajan LEAP robot and passed through a custom-prepared immobilized pepsin column (2.1 × 50 mm) at 0.2 mL min^-1^ using an Agilent 1290 LC system. The resulting peptic peptides were trapped and desalted on a ZORBAX 300SB-C8 trap column (2.1 × 12.5 mm) and separated on an XSelect CSH C18 column (2.1 × 50 mm, 2.5 µm; Waters) using a linear 14-min gradient from 5% to 100% solvent B. Solvent A was 100% H_2_O containing 0.1% formic acid, and solvent B was 80:20 acetonitrile:H_2_O containing 0.1% formic acid. Peptide *m/z* values were measured on a Bruker Maxis II Q-TOF ETD mass spectrometer. Raw data were analyzed using HDExaminer v3.4.1, exported directly from HDExaminer, and submitted to Kingfisher^57^ for downstream statistical analysis and visualization (see Supplementary Methods).

### Native and denaturing mass spectrometry

Native MS was performed under soft ionization and transmission conditions optimized to preserve noncovalent CK-BB dimers^89,90^. Spray voltages were maintained between 0.6 and 1.2 kV, and collision energies were adjusted as needed. Denaturing ESI-MS was performed on a Bruker MaXis II Q-TOF mass spectrometer. Samples were trapped on a ZORBAX 300SB-C3 column, desalted with water/acetonitrile/formic acid and eluted with a short acetonitrile gradient. Protein mass spectra were deconvoluted using Intact or UniDec.

### Molecular dynamics simulations

All-atom MD simulations were performed for four CK-BB states: apo pH 5.5, apo pH 7.3, ligand-bound pH 5.5 and ligand-bound pH 7.3. Starting structures were derived from AlphaFold3 models^56^. For the pH 5.5 systems, protonation states were assigned using PROPKA^91^ and applied symmetrically to both protomers. Protonated residues included Asp62, Glu19, Glu80, Glu155, Glu231, His0 and His66. No additional protonation was applied to the pH 7.3 systems. Systems were prepared in explicit solvent using CHARMM-GUI^92^, solvated with TIP3P water^93^, neutralized with Na^+^ and Cl^−^ ions, and supplemented with 0.15 M NaCl. The CHARMM36m force field was used^94^. Each solvated system contained approximately 179,000 atoms.

Initial energy minimization was performed for 10,000 conjugate-gradient steps. Systems were then equilibrated for approximately 19 ns in NAMD^95^ using the standard CHARMM-GUI restrained protocol in the NVT ensemble at 310 K, with a Langevin thermostat damping coefficient of 1.0 ps^-1^. A subsequent 15-ns pre-production run was performed in the NPT ensemble at 310 K and 1 atm, with pressure maintained using the Nosé–Hoover Langevin piston method^96^ and a 2-fs integration step. Equilibrated systems were then converted to Anton-compatible format, and triplicate 1-µs production simulations were performed on the Anton 3 supercomputer^97^ in the NPT ensemble using a Desmond-compatible simulation framework. Temperature was maintained at 310 K using an antithetic thermostat with an interval of 48 timesteps, and pressure was maintained isotropically at 1.0 bar using a Monte Carlo barostat with an interval of 480 timesteps. A 2.5-fs integration step with RESPA multiple time stepping was used. Trajectories were analyzed for RMSD, RMSF, interdomain angles and residue-specific dynamics using custom scripts and VMD^98^. RMSD was calculated relative to the first frame of each production trajectory after least-squares superposition of protein Cα atoms. For angle analysis, the relative orientation between equivalent regions of adjacent protomers was quantified from MD trajectories using custom Tcl scripts in VMD. Each frame was aligned to the initial structure by least-squares fitting of protein Cα atoms to remove global translation and rotation. For each protomer, the region of interest was defined by Cα atoms from residues 216–220, 225–229 and 235–242, and its orientation was defined by the principal axis of the Cα inertia tensor corresponding to maximal spatial extension. The inter-protomer angle was calculated from the dot product of the normalized principal axes and converted to degrees. To remove sign ambiguity, angles greater than 90° were mapped to their supplementary values, yielding a 0–90° measure of relative domain opening and closure between protomers.

## Data availability

The generated data, including data from DEER, HDX-MS and native mass spectrometry experiments, are available in the manuscript or Supplementary Materials. Source data are provided with this paper. Additional datasets, including raw DEER data, MD input files, initial and final coordinates, analysis scripts, analysis data, DEER-guided AlphaFold models and refined models, have been deposited in the Zenodo repository maintained by CERN: https://doi.org/10.5281/zenodo.21539483. Other data supporting this study are available from the corresponding author upon request.

## Acknowledgments

This work was supported by NIH grant R37-CA265877 to R.D. The molecular dynamics studies were supported by NIH grant R35-GM147423 to M.M. Anton 3 computer time was provided by the Pittsburgh Supercomputing Center through NIH grant R24-GM154042. The Anton 3 system at PSC is made available by D. E. Shaw Research. The membrane binding and curvature sensing studies were supported by NIH grant R35-GM147333 to W.F.Z.

## Author contributions

R.D. designed the experiments with input from all authors and supervised the study. S.G. prepared constructs, expressed and purified WT and mutant proteins, and performed biochemical validation. R.D. and S.G. performed the EPR experiments and analyzed the data. N.K.M. performed native MS and HDX-MS experiments and analyses. M.L.G. supervised the MS analyses. O.H.K. and W.F.Z. performed and supervised the membrane binding experiments and analyses. A.S. and M.M. performed and supervised the molecular dynamics simulations and analyses. C.K., S.K., A.U. and D.W.P. performed and supervised the cell-imaging experiments. T.W. and R.D. performed the structural modeling and analyses. I.M. contributed to protein preparation. R.D. wrote the manuscript with input from all authors. N.K.M. and O.H.K. contributed equally to this work, as did C.K. and A.S.

## Supplementary Information

This PDF file includes

Supplementary Methods

Supplementary Figs. 1-17

Supplementary Table 1

## Supplementary Methods

### Vesicle preparation

1-palmitoyl-2-oleoyl-glycero-3-phosphocholine (POPC), 1-palmitoyl-2-oleoyl-glycero-3-phospho-L-serine (sodium salt) (POPS), 1,2-dioleoyl-sn-glycero-3-phosphoethanolamine (DOPE), 1-stearoyl-2-arachidonoyl-sn-glycero-3-phosphoinositol (ammonium salt) (SAPI), and cholesterol (plant) were purchased from Avanti Research. ATTO 647N-labeled 1,2-dipalmitoyl-sn-glycero-3-phosphoethanolamine (ATTO 647N-DPPE) was purchased from Leica Microsystems, and dipalmitoyl-decaethylene glycol-biotin (DP-EG15-biotin) was provided by D. Sasaki (Sandia National Laboratories, Livermore, CA)^1^.

Lipid aliquots were combined at the following molar percentages for tethered-vesicle assays: 13.5% POPC, 34.5% DOPE, 22% POPS, 9% SAPI, 20% cholesterol, 0.5% DP-EG15-biotin, and 0.5% ATTO 647N-DPPE for the plasma membrane mimetic; and 24.5% POPC, 24.5% DOPE, 10% POPS, 40% cholesterol, 0.5% DP-EG15-biotin, and 0.5% ATTO 647N-DPPE for the synaptic vesicle mimetic. Lipid mixtures were dried under a gentle stream of nitrogen to remove chloroform and placed under vacuum for ≥2 h. Dried lipid films were rehydrated in 20 mM sodium phosphate and 100 mM NaCl, pH 7.3 (Neutral Buffer), and subjected to five freeze–thaw cycles by immersion in liquid nitrogen for 1 min followed by thawing at 50 °C for 5 min. Vesicles were then extruded through 50-nm polycarbonate membranes (Sterlitech). Plasma membrane-mimetic vesicles were handled under nitrogen to minimize oxidation of the polyunsaturated SAPI lipid.

### Slide passivation and vesicle tethering

Biotin-PEG-silane (5 kDa) and mPEG-silane (5 kDa) were purchased from Laysan Bio. Glass slides were cleaned and passivated as described previously^2–6^. Briefly, slides were bath-sonicated in 70% ethanol, rinsed, and sequentially incubated in concentrated KOH/peroxide and HCl/peroxide solutions at 80 °C for 10 min each, followed by rinsing with water and drying. Cleaned slides were passivated with 55 μL of PEG-silane solution containing biotin-PEG-silane and mPEG-silane in isopropanol, with glacial acetic acid added to catalyze silane coupling to surface hydroxyl groups, and incubated at 50 °C for 1 h. Silicone gaskets (Grace Bio-Labs) containing 5-mm-diameter wells were applied to the passivated slides. Each well was incubated with 30 μL of 3.33 μM NeutrAvidin for 20 min and rinsed with Neutral Buffer to remove unbound NeutrAvidin. Vesicles were then added at a lipid concentration of 1 μM, incubated for 10 min, and rinsed with Neutral Buffer to remove unbound vesicles. For acidic conditions, wells were subsequently exchanged into 20 mM sodium phosphate and 100 mM NaCl, pH 5.5. CK-BB, with or without substrates, was added at the indicated concentrations and incubated for 1 h before imaging.

### Confocal microscopy

Fluorescence imaging was performed on a Leica Stellaris 5 laser-scanning confocal microscope equipped with a Leica HC PL APO 63x oil-immersion objective (NA 1.4). AZDye 488-labeled CK-BB was excited at 488 nm, with emission collected from 493–558 nm; ATTO 647N-labeled vesicles were excited at 638 nm, with emission collected from 643–749 nm. The optical zoom was adjusted to yield a pixel size of 70 x 70 nm, and images were acquired at a line-scanning frequency of 400 Hz. For fluorescence recovery after photobleaching (FRAP), three pre-bleach images were acquired over 2 s, followed by photobleaching of the region of interest at 100% laser power for five frames (∼4 s). Fluorescence recovery was then monitored by acquiring 50 post-bleach images at 5-s intervals.

### Image processing and calibration

Image analysis was performed as described previously^2–4^. Diffraction-limited fluorescent puncta corresponding to vesicles and membrane-bound protein were identified and tracked using cmeAnalysis. Vesicles were identified in the lipid channel and retained only when detected in three consecutive frames. Vesicles with diameters >250 nm were excluded to restrict the analysis to diffraction-limited particles. CK-BB binding to individual vesicles was quantified from colocalized fluorescence intensity in the protein channel. Vesicle fluorescence intensity was converted to diameter by determining the fluorescence-intensity distribution for each vesicle population and deriving a scaling factor by matching the square root of this distribution to the corresponding diameter distribution measured by dynamic light scattering (DLS). For curvature-dependent FRAP analyses of the nominal 50-, 100-, and 150-nm vesicle populations, vesicles were grouped into diameter ranges of 40–60, 90–110, and 140–160 nm, respectively. Single-molecule fluorescence intensity was calibrated by adsorbing AZDye 488-labeled CK-BB onto glass coverslips and measuring the resulting punctate fluorescence-intensity distribution. The peak of this distribution was assigned as the fluorescence intensity of a single labeled CK-BB molecule and used to determine the number of membrane-bound proteins per vesicle. Calibration values were adjusted as necessary to account for differences in detector gain between imaging experiments.

### FRAP analysis

The mobile fraction (*x_m_*) and apparent dissociation rate constant (*k_off_*) of membrane-bound CK-BB were determined by fitting fluorescence-recovery curves to Equation (1) as described previously^3^:

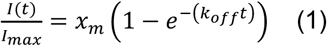

where *I*(*t*) is the average fluorescence intensity within the bleached region at time *t* after photobleaching, *I_max_* is the average fluorescence intensity before photobleaching, *x_m_* is the mobile fraction of membrane-bound CK-BB, and *k_off_* is the apparent dissociation rate constant of the mobile fraction.

### pH correction and HDX calculations

Because intrinsic backbone amide exchange slows at lower pH in the base-catalyzed regime, labeling times were adjusted to separate intrinsic pH effects on exchange chemistry from condition-dependent changes in protein dynamics or solvent exposure. The pH correction factor was calculated relative to the reference condition, pH 7.3, using Equation (2):

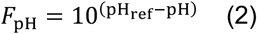

where pH_ref_ = 7.3. A reference labeling time at pH 7.3 was converted to the matched lower-pH labeling time using Equation (3):

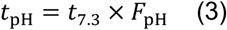

This procedure yielded correction factors of 6.3, 20.0 and 63.1 at pH 6.5, 6.0 and 5.5, respectively (Supplementary Table 1). These extended labeling times allowed deuterium uptake to be compared across pH conditions on matched intrinsic-exchange windows. Peptide centroid masses were calculated from the isotope envelope using Equation (4):

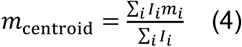

where *I_i_* and *m_i_* are the intensity and mass of isotope peak *i*. Deuterium uptake was calculated from the centroid mass after labeling (*m_t_*), the undeuterated centroid (*m*_0_) and the fully deuterated or maximum-exchange reference (*m*_100_) using Equation (5):

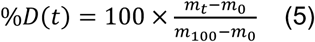

Pairwise deuterium differences were calculated for each comparison using Equation (6):

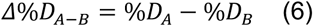

Positive values indicated greater exchange in condition A relative to condition B. Together, the pH-adjusted exchange times, centroid-based uptake calculations, and peptide-level comparisons enabled condition-dependent changes in local backbone dynamics and/or solvent exposure to be distinguished from intrinsic pH-dependent changes in amide exchange kinetics.

### Peptide identification and HDX data analysis

Peptide identification was performed by LC-MS/MS in data-dependent acquisition mode on a Bruker Maxis II Q-TOF ETD mass spectrometer prior to conducting HDX. MS/MS spectra were searched and curated in Protein Metrics software using a score cutoff of 200 and manual validation of fragment assignments. HDX data were analyzed with Kingfisher^7^ using a hybrid significance-testing approach that combines a peptide-level statistical test with a global experimental-error threshold^8,9^. All peptide isotope envelopes used in the differential HDX-MS analysis were manually inspected for EX1-type bimodal isotope distributions, and no evidence of EX1 exchange was observed.

For each pairwise comparison, Welch’s *t* statistic was calculated using Equation (7):

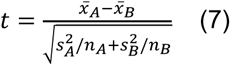

with Welch-Satterthwaite degrees of freedom calculated using Equation (8):

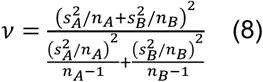

To estimate the magnitude threshold required for significance, a pooled standard deviation was calculated across replicate measurements using Equation (9):

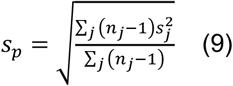

The propagated standard error for a difference between two states was then calculated using Equation (10):

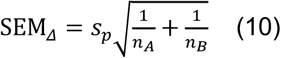

where *n_A_* and *n_B_* denote the numbers of experimental replicates for protein states A and B, respectively. The global significance threshold was calculated using Equation (11):

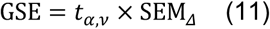

where *t*_α_,_ν_ is the critical value from Student’s *t*-distribution, selected according to the predefined significance level (α), and the applicable degrees of freedom (ν). A peptide was considered significantly changed only when the absolute deuterium difference exceeded the global significance threshold, and the Welch *t* test met the indicated p-value cutoff. Peptides meeting *p* < 0.001 were displayed as stringent hits, peptides meeting *p* < 0.05 were displayed as less stringent hits, and peptides failing either criterion were considered not significantly different. A peptide was classified as significant if at least one time point satisfied both criteria.

---

**Supplementary Figure 1.**
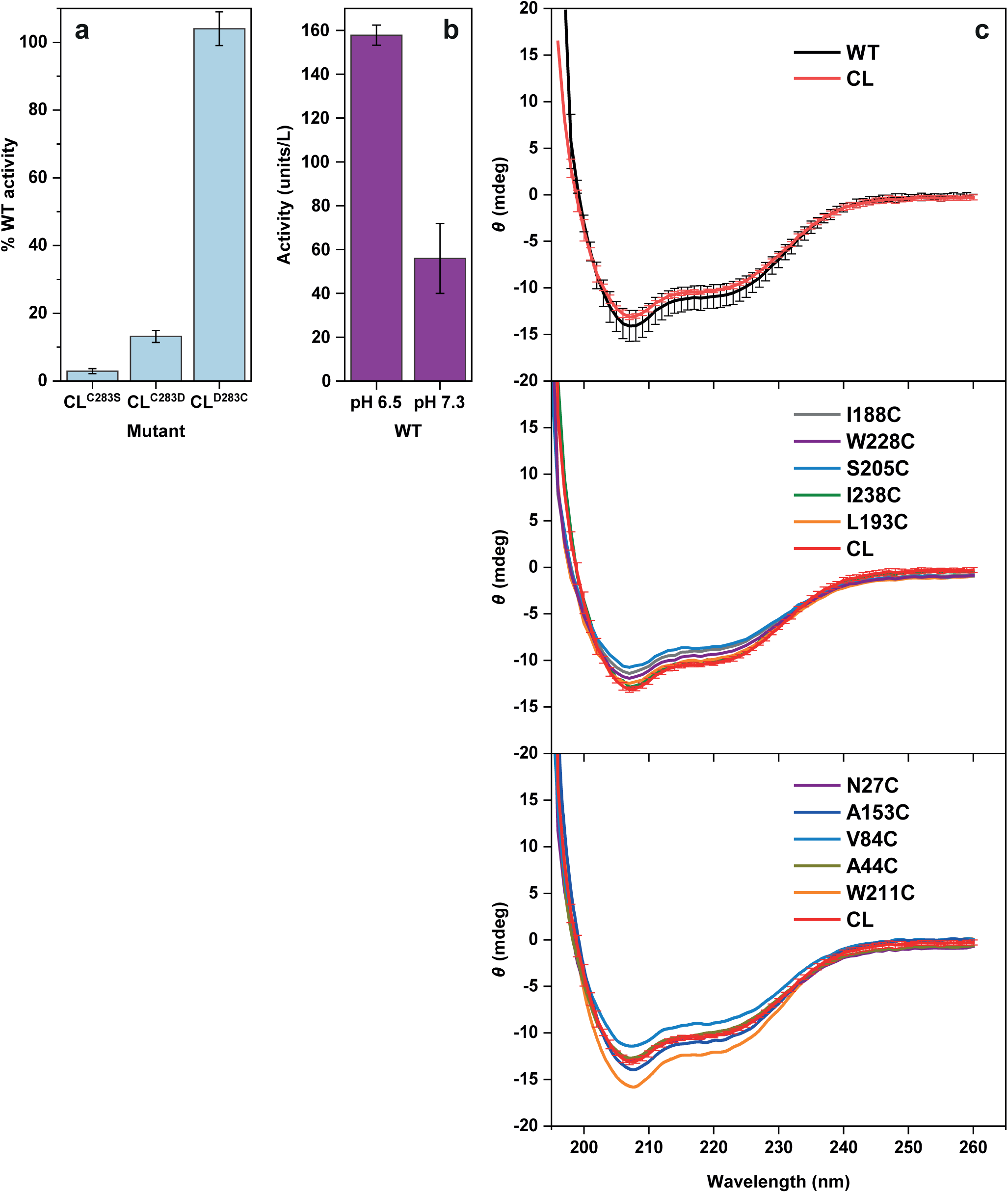
Functional and structural integrity analyses. (**a**) Creatine kinase activity of cysteine-less (CL) CK-BB variants normalized to WT. Mutation of the catalytic Cys283 to serine strongly reduced activity, whereas the CL^C283D^ background retained measurable activity and was used for DEER analyses. Mutation of the remaining native cysteines did not affect catalytic activity, as shown by the CL^D283C^ construct. Each assay was performed in triplicate (technical replicates; *n* = 3). (**b**) WT CK-BB activity is enhanced at mildly acidic pH relative to pH 7.3. (**c**) Far-UV circular dichroism spectra of WT, CL, and spin-labeled CK-BB mutants show closely similar profiles, indicating that cysteine substitution and spin labeling do not subs1tantially perturb global secondary structure. CD measurements for WT and CL were performed in triplicate. Values represent mean ± s.d. Source data are provided as a Source Data file.

**Supplementary Figure 2.**
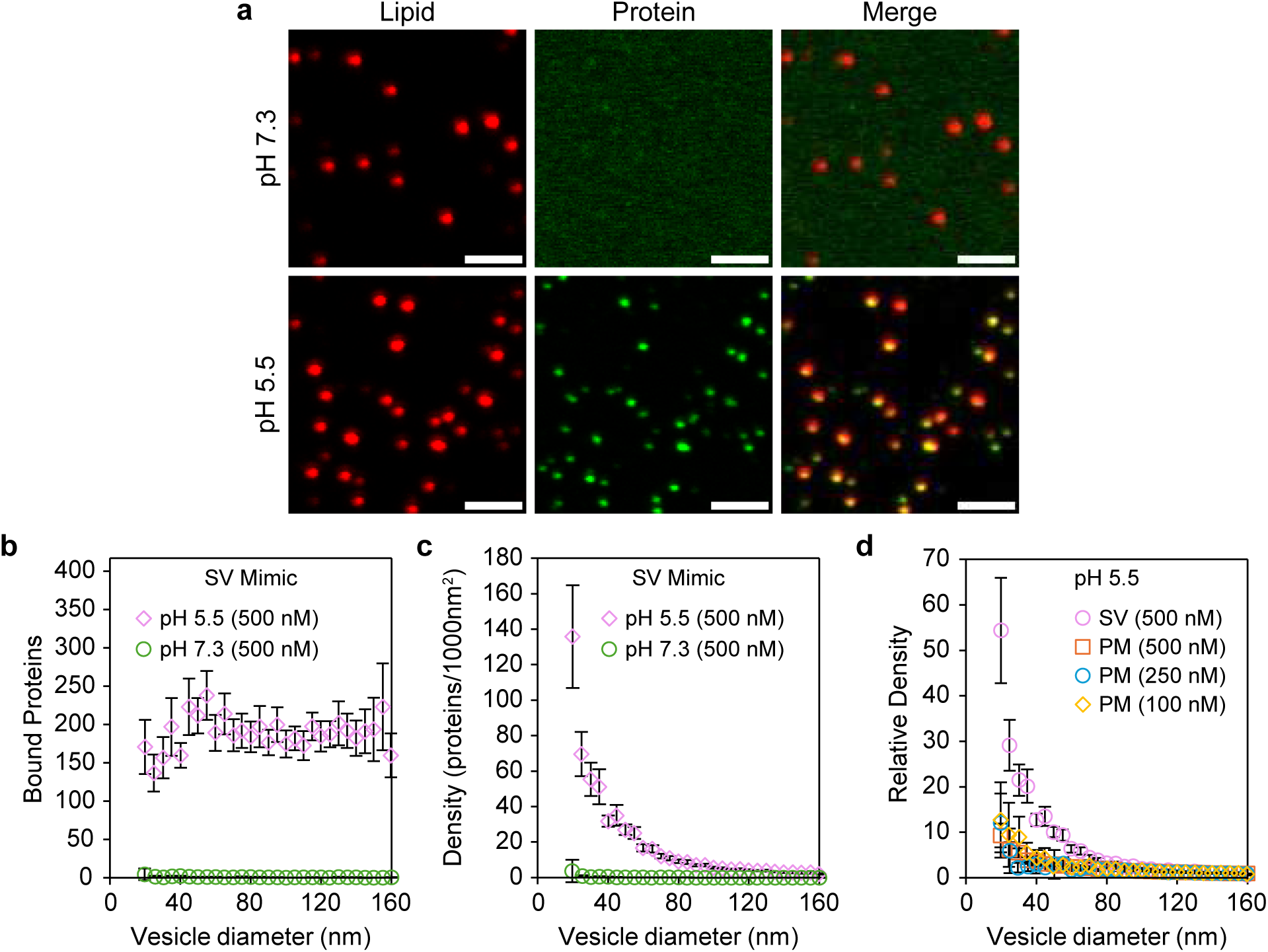
Acidification promotes curvature-sensitive CK-BB association with synaptic vesicle mimetics. (**a**) Representative lipid, CK-BB and merged fluorescence micrographs of tethered synaptic vesicle (SV)-mimetic vesicles incubated with apo CK-BB at pH 7.3 or 5.5. (**b**) Number of membrane-bound apo CK-BB molecules as a function of vesicle diameter. (**c**) Membrane-bound CK-BB density as a function of vesicle diameter. (**d**) Relative CK-BB density normalized to the mean density on vesicles 130–160 nm in diameter. PM-mimetic data from Fig. 2g are shown for comparison. For **b**–**d**, vesicles 20–160 nm in diameter were grouped into 29 bins at 5-nm intervals, and each point represents the mean within the corresponding bin. Numbers of imaged and processed vesicles were *N* = 8296 (pH 5.5, 500 nM) and *N* = 2474 (pH 7.3, 500 nM). Data for pH 5.5, 500 nM were pooled across three biological replicates (*n* = 3), whereas pH 7.3, 500 nM represents an individual measurement (*n* = 1). Error bars indicate 95% confidence intervals of the mean. Scale bars, 2 μm.

**Supplementary Figure 3.**
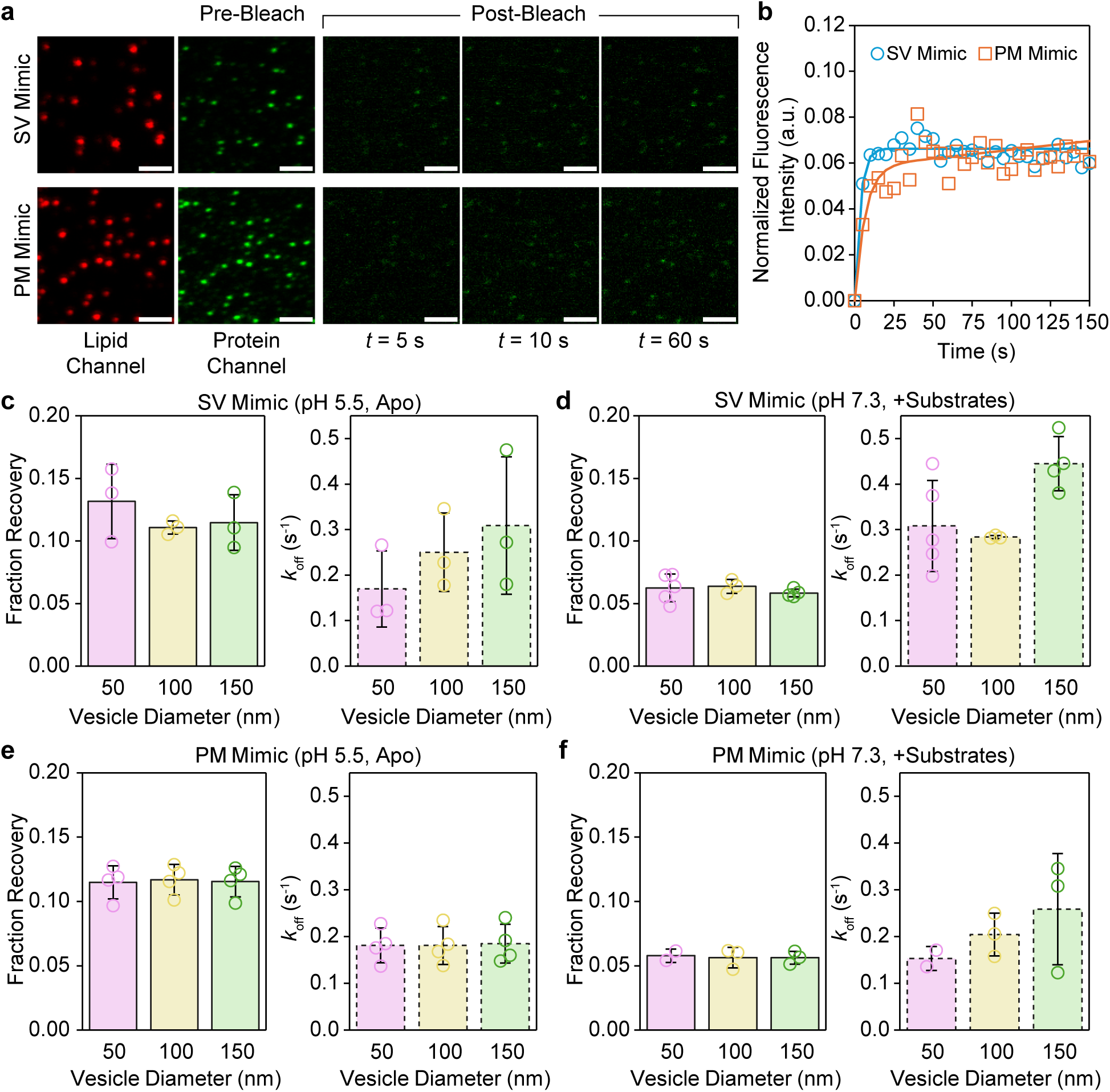
FRAP reveals persistent, slowly exchanging CK-BB membrane association. (**a**) Representative fluorescence micrographs of membrane-bound CK-BB and lipids before photobleaching and during fluorescence recovery. (**b**) Representative fluorescence recovery curves fitted using Equation 1. Representative images and recovery curves in **a**,**b** are shown for substrate-bound CK-BB at pH 7.3. (**c**–**f**) Recovered protein fraction and corresponding apparent dissociation rate constants (*k*_off_) for 1 μM apo CK-BB at pH 5.5 and substrate-bound CK-BB at pH 7.3 on synaptic vesicle (SV)- and plasma membrane (PM)-mimetic vesicles. Data are mean ± s.d. from 2–5 independent FRAP measurements per condition. Scale bars, 2 μm.

**Supplementary Figure 4.**
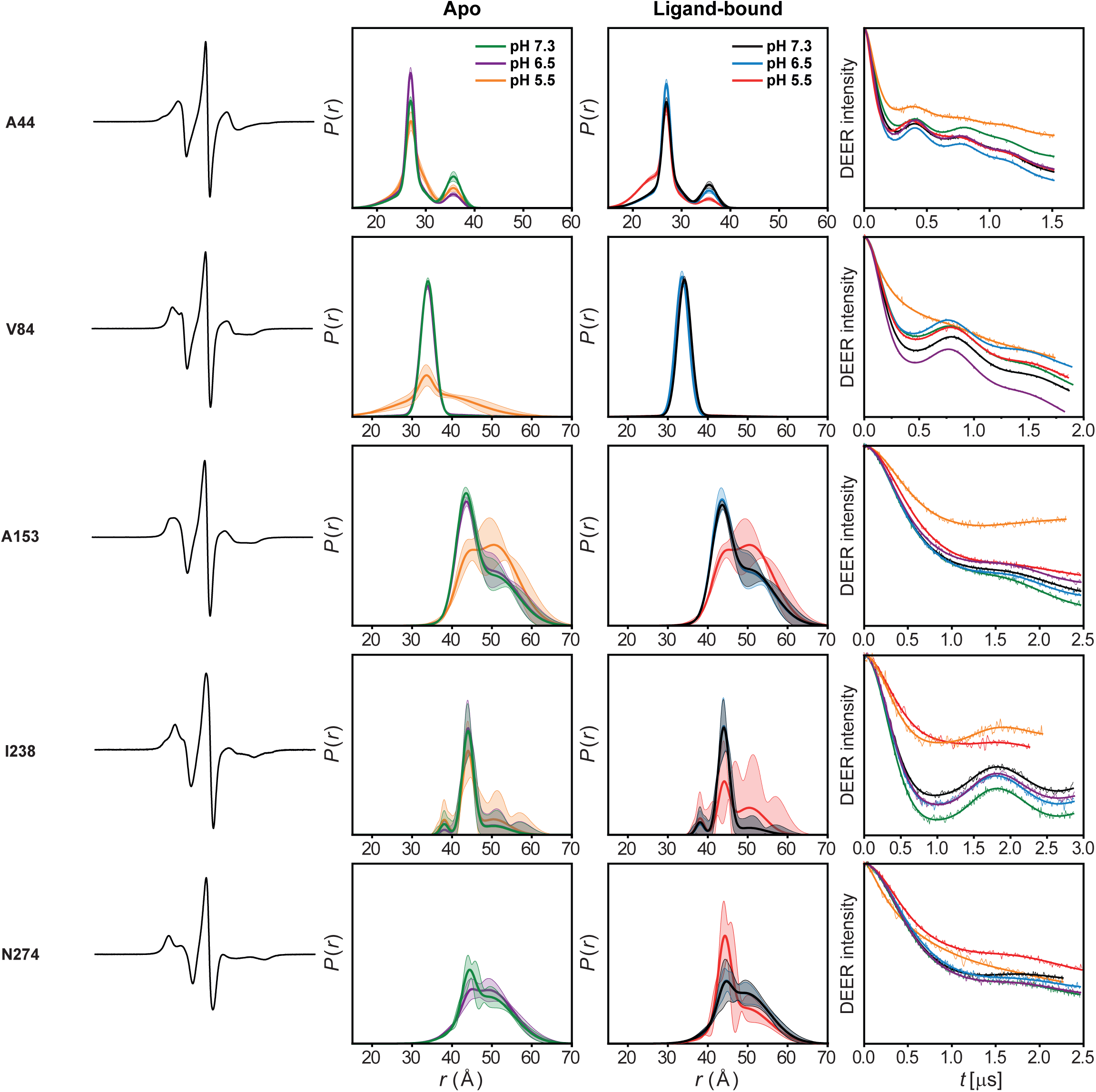
DEER data analysis for inter-protomer spin-label pairs across the convex side of the NTD dimer interface and C-terminal beta-sheet core. For each mutant, from left to right, CW-EPR spectrum, DEER-derived distance distributions and primary DEER traces with corresponding fits are shown for apo and ligand-bound CK-BB at physiological and acidic pH. These reporters correspond to the convex-side measurements summarized in Fig. 3 and include sites spanning the NTD dimer interface and CTD beta-sheet core. Confidence bands indicate 2σ uncertainty from fitting of the primary DEER traces.

**Supplementary Figure 5.**
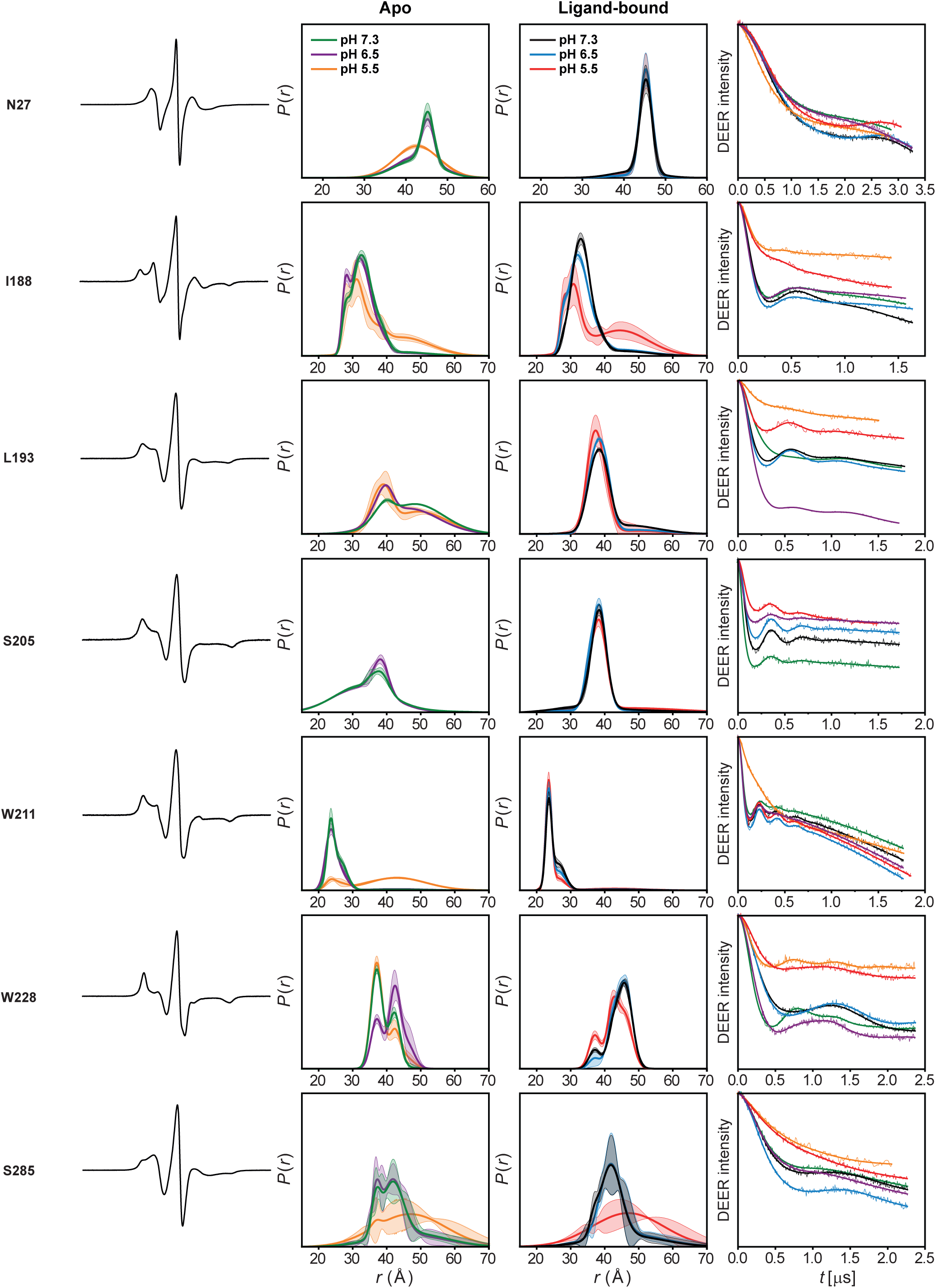
DEER data analysis for inter-protomer spin-label pairs across the concave side of the CK-BB dimer. For each mutant, from left to right, CW EPR spectrum, DEER-derived distance distributions and primary DEER traces with corresponding fits are shown for apo and ligand-bound CK-BB at physiological and acidic pH. These reporters correspond to the concave-side measurements summarized in Fig. 4 and span the Ser199-containing region and neighboring active-site elements. Confidence bands indicate 2σ uncertainty from fitting of the primary DEER traces.

**Supplementary Figure 6.**
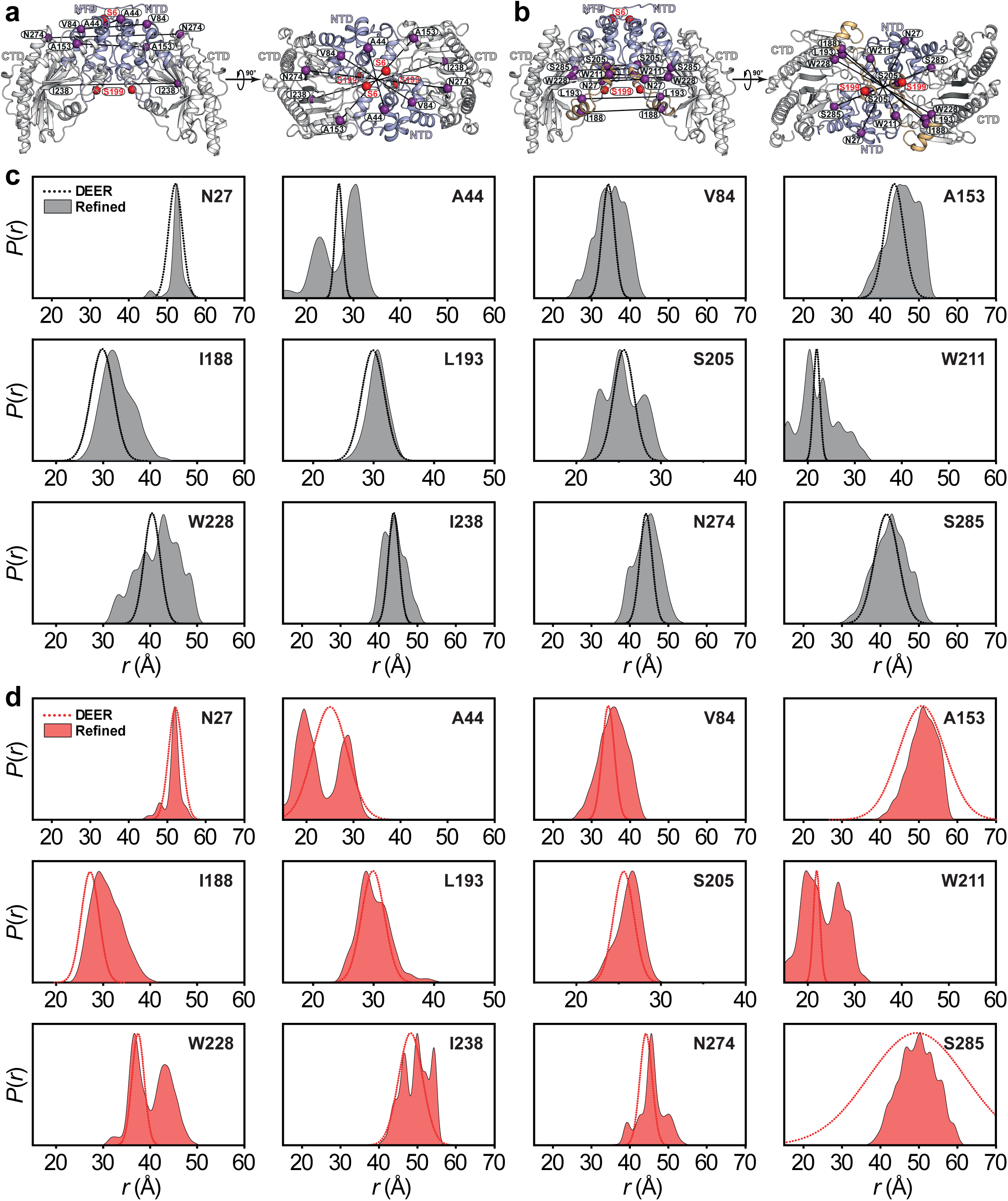
DEER-guided refinement of ligand-bound CK-BB at physiological and acidic pH. (**a**,**b**) Inter-protomer spin-label pairs used for DEER measurements across the convex (**a**) and concave (**b**) sides of the CK-BB dimer. Spin-label sites are shown as purple spheres, and Ser6 and Ser199 phosphorylation sites are shown as red spheres. (**c**,**d**) Predicted distance distributions from DEER-refined ligand-bound CK-BB models at pH 7.3 (**c**, gray) and pH 5.5 (**d**, red), compared with the corresponding experimental DEER distance subpopulations shown as dotted lines. The refined models closely recapitulate the experimental distance subpopulations.

**Supplementary Figure 7.**
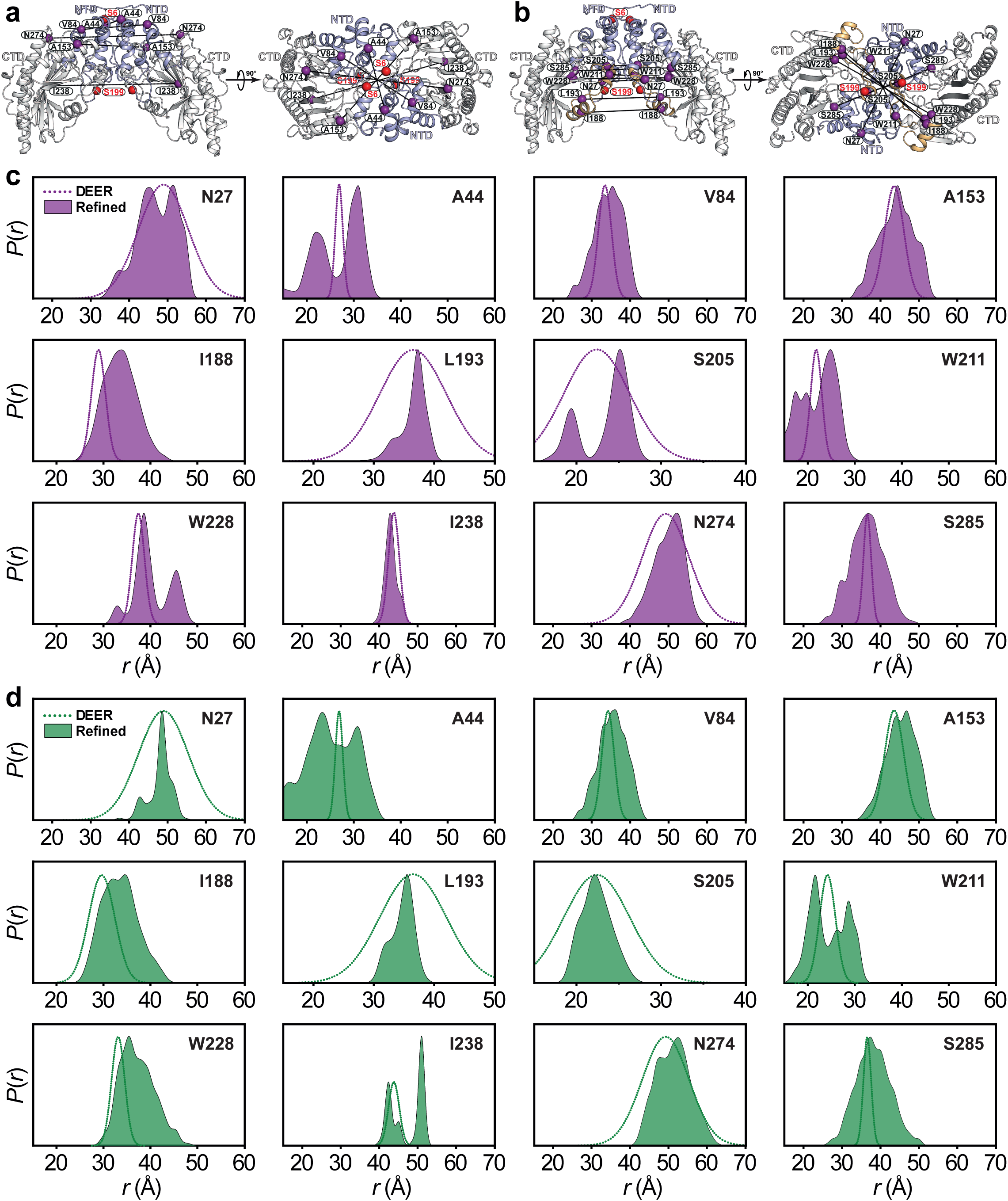
DEER-guided refinement of apo CK-BB at physiological and acidic pH. (**a**,**b**) Inter-protomer spin-label pairs used for DEER measurements across the convex (**a**) and concave (**b**) sides of the CK-BB dimer. Spin-label sites are shown as purple spheres, and Ser6 and Ser199 phosphorylation sites are shown as red spheres. (**c**,**d**) Predicted distance distributions from DEER-refined apo CK-BB models at pH 6.5 (**c**, purple) and pH 7.3 (**d**, green), compared with the corresponding experimental DEER distance subpopulations shown as dotted lines. The refined apo models closely recapitulate the experimental distance subpopulations.

**Supplementary Figure 8.**
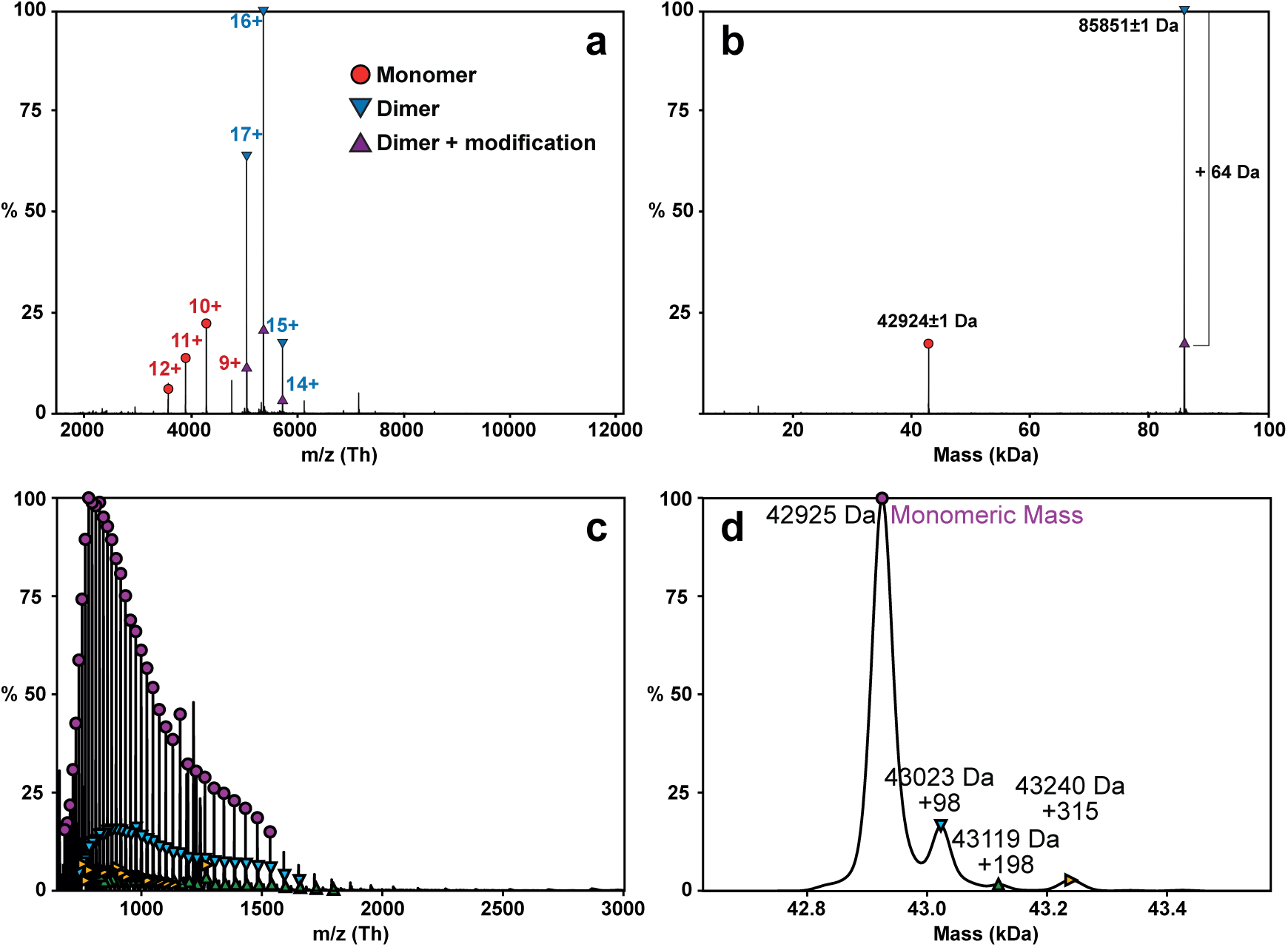
Native MS confirms stable dimer formation by CK-BB. **(a,b)** Raw (**a**) and deconvoluted (**b**) native mass spectra of CK-BB. Charge-state series are labeled and color-coded for monomeric and dimeric species, with matching symbols linking raw and deconvoluted peaks. The dominant dimer peak is observed at 85,851 ± 1 Da, whereas the monomeric species likely reflects partial dissociation of the native dimer during ionization. A minor +64 Da species indicates an unidentified post-translational or chemical modification. (**c**,**d**) Raw (**c**) and deconvoluted (**d**) denaturing MS spectra showing a monomeric mass of 42,925 Da, in close agreement with the theoretical CK-BB mass of 42,925.5 Da; minor higher-mass species are also observed.

**Supplementary Figure 9.**
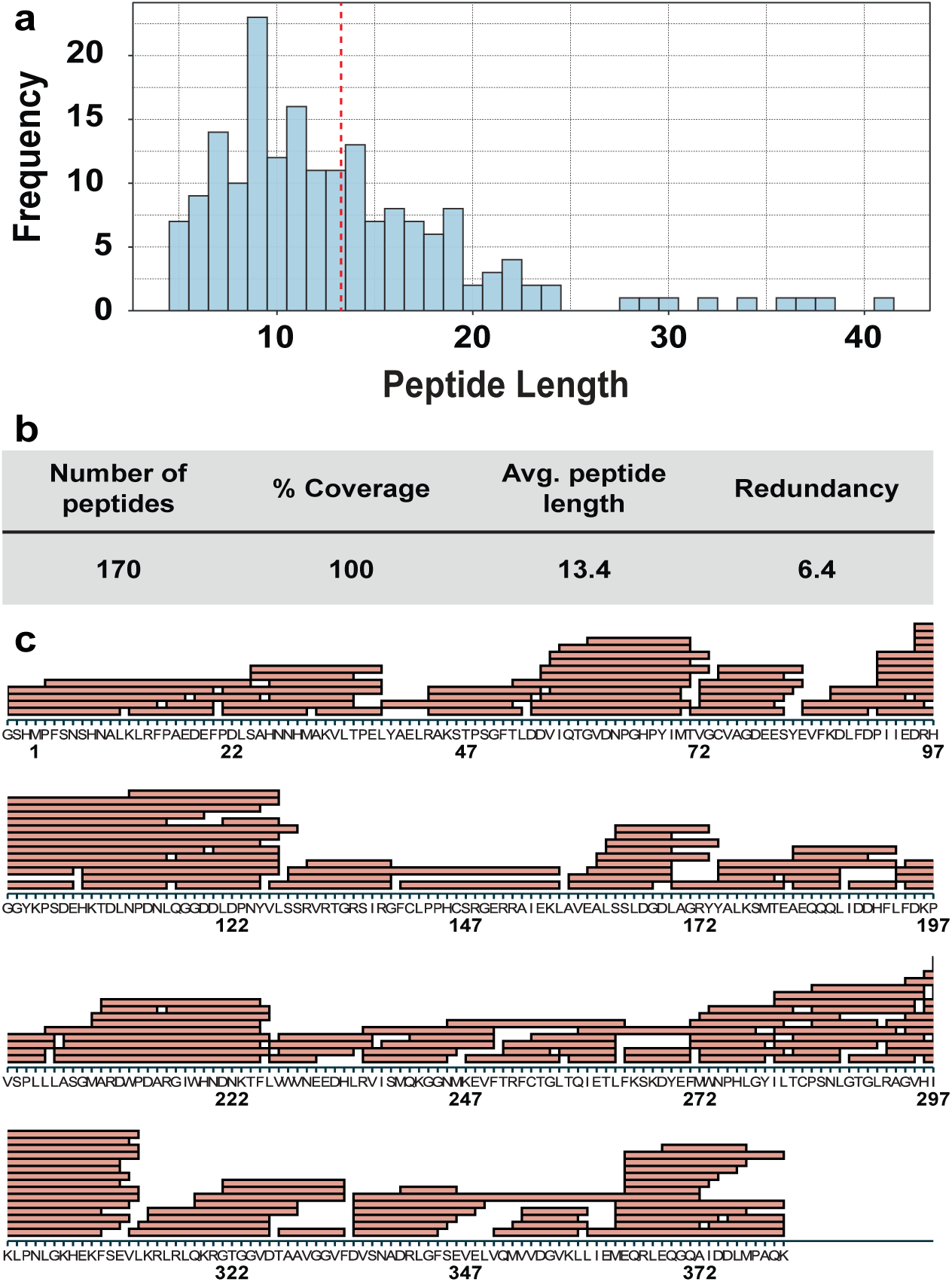
HDX-MS peptide coverage and digestion quality for CK-BB. (**a**) Frequency distribution of identified peptides by length, with the dashed line indicating the average peptide length. (**b**) Summary of HDX-MS coverage metrics, including total number of peptides, sequence coverage, average peptide length and redundancy. (**c**) Peptide coverage map of CK-BB, with each red bar representing an MS/MS-verified peptide and its position along the protein sequence. The dataset provides complete sequence coverage with high redundancy, supporting robust peptide-level HDX analysis. Figures were generated using Kingfisher software.

**Supplementary Figure 10.**
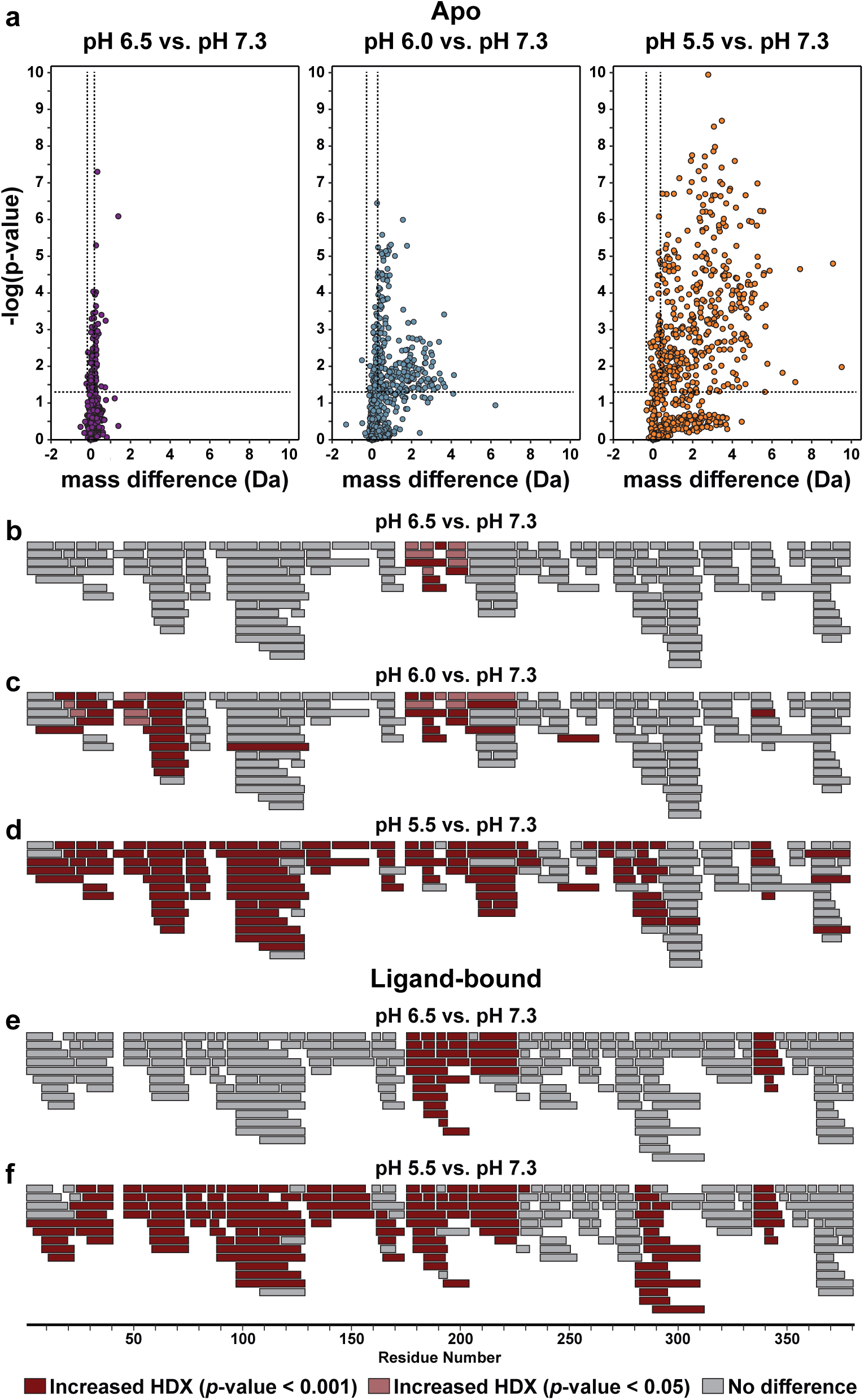
HDX-MS reveals enhanced CK-BB backbone dynamics under acidic conditions. (**a**) Volcano plots showing pH-dependent changes in deuterium uptake for apo CK-BB relative to pH 7.3. The vertical axis shows statistical significance (−log_10_ *p* value from *t* tests), with higher values indicating more significant changes; data points above the dotted threshold lines were considered significant. Lower pH progressively increases both the magnitude and significance of deuterium uptake, indicating enhanced conformational dynamics. (**b**–**f**) Peptide coverage maps for apo (**b**–**d**) and ligand-bound (**e**,**f**) CK-BB, showing regions with significantly increased HDX at the indicated pH values relative to pH 7.3. Dark red and light red indicate peptides meeting stringent (*p* < 0.001) and less stringent (*p* < 0.05) significance thresholds, respectively; gray indicates peptides that failed the hybrid significance test, either by not exceeding the global error threshold or by not meeting the minimum *p*-value cutoff.

**Supplementary Figure 11.**
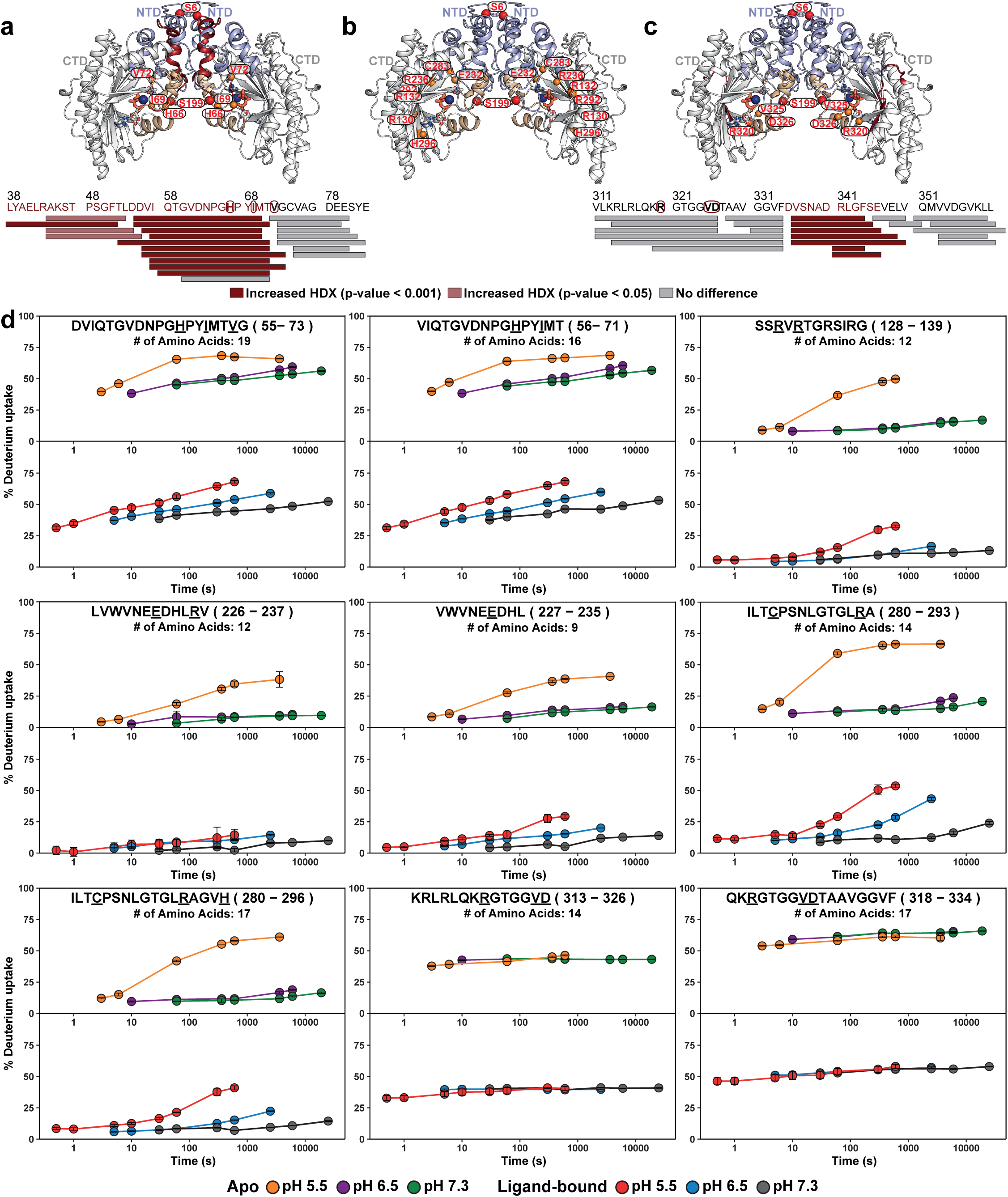
Functional CK-BB regions show ligand-sensitized acid-dependent exchange. (**a**) CK-BB model highlighting the NTD interface and adjacent conserved flexible loop, including His66, Ile69 and Val72, which show increased deuterium uptake under acidic conditions. Functional residues in this region are shown as orange spheres. (**b**) Additional catalytic and substrate-binding residues on the concave surface. (**c**) CK-BB model highlighting a C-terminal beta-sheet core region that shows increased deuterium uptake in the ligand-bound state under acidic conditions. (**d**) HDX uptake plots for peptides spanning catalytic and substrate-binding regions. In apo CK-BB, peptides containing His66, Ile69, Val72, Arg130, Arg132, Glu232, Arg236, Cys283, Arg292 and His296 show little difference between pH 7.3 and 6.5, but markedly increased exchange at pH 5.5, consistent with a threshold-like acid-induced increase in local backbone dynamics and/or solvent exposure. In the ligand-bound state, most of these functional regions, including peptides containing His66, Ile69, Val72, Glu232, Cys283, Arg292 and His296, display graded pH-dependent increases in exchange, consistent with substrate-dependent sensitization of local dynamics. Global significance threshold, 0.32 Da; *p* < 0.001.

**Supplementary Figure 12.**
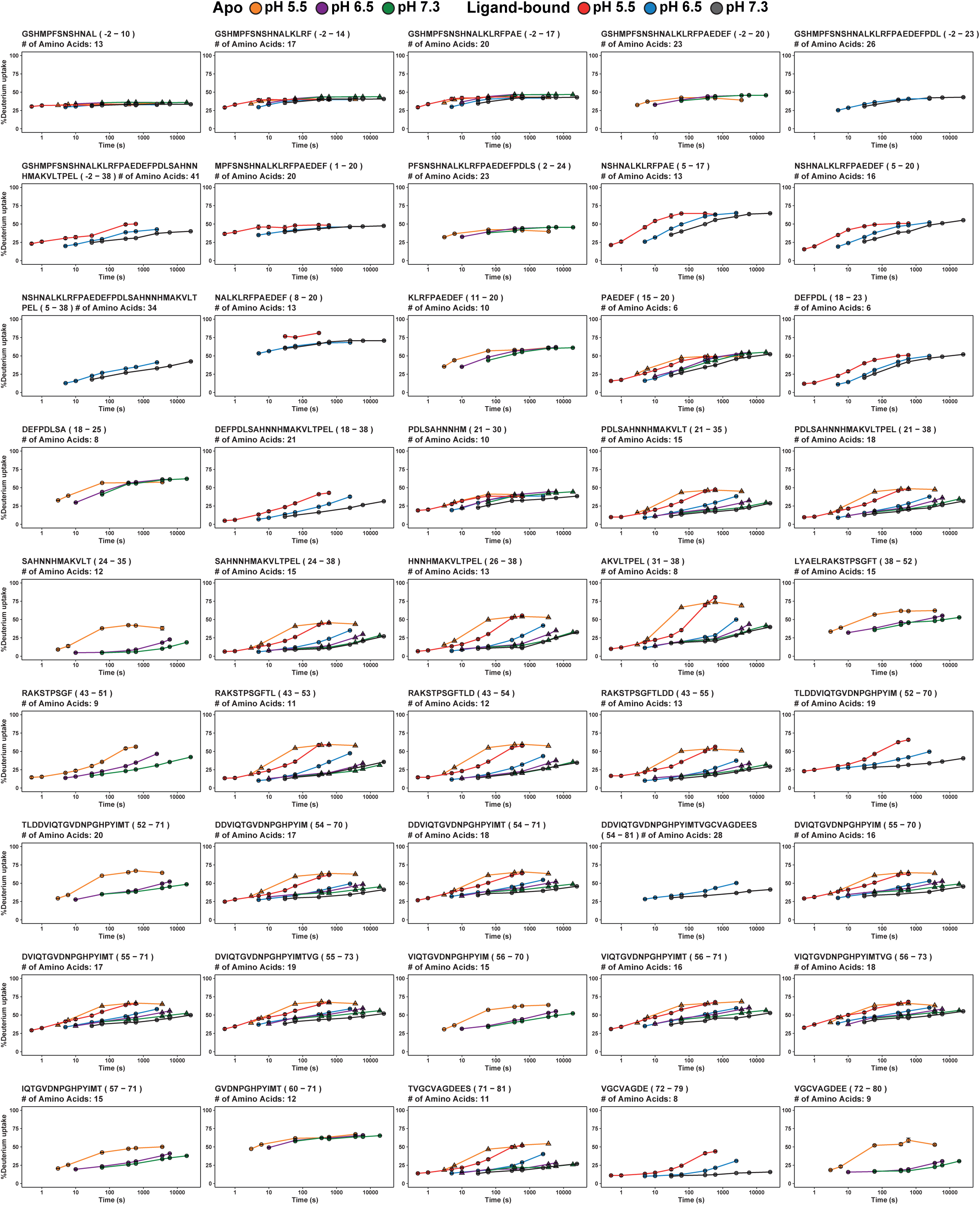
HDX uptake kinetics across the CK-BB N-terminal region and flexible active-site loop. HDX uptake plots for overlapping peptides spanning residues −2 to 81, shown as percentage deuterium uptake over exchange time for apo and ligand-bound CK-BB at pH 7.3, 6.5 and 5.5. Peptides in the extreme N-terminal region show relatively modest pH- and ligand-dependent differences, whereas peptides spanning residues ∼24–38 and the conserved flexible loop around residues ∼52–73 exhibit pronounced acid-dependent increases in exchange. Several peptides containing the His66/Ile69 region show a threshold-like increase in the apo state, with stronger exchange at pH 5.5, while ligand-bound CK-BB displays more graded pH-dependent exchange, consistent with substrate-dependent sensitization of local backbone dynamics and/or solvent exposure.

**Supplementary Figure 13.**
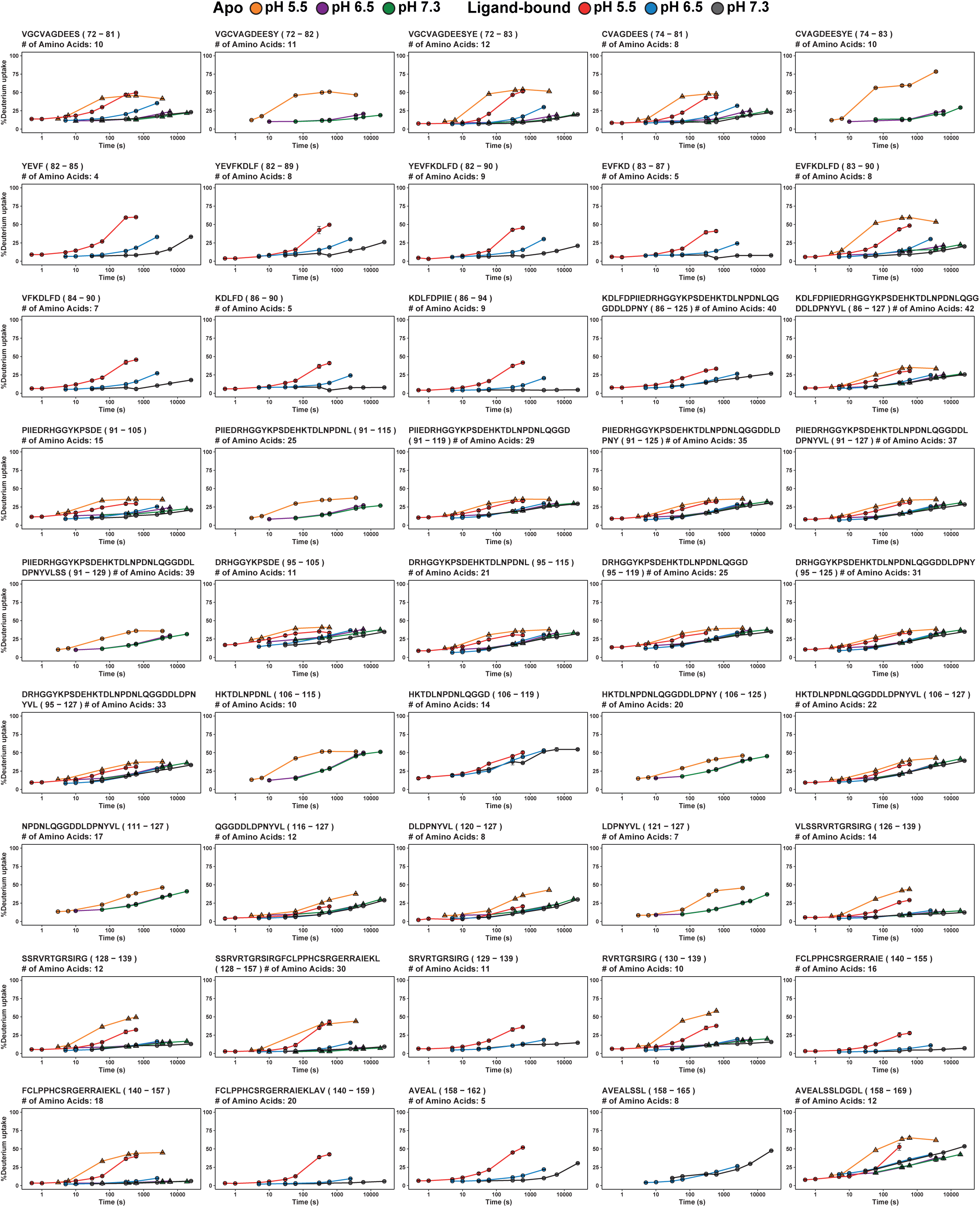
HDX uptake kinetics across CK-BB residues 72–169. HDX uptake plots for overlapping peptides spanning residues 72–169, shown as percentage deuterium uptake over exchange time for apo and ligand-bound CK-BB at pH 7.3, 6.5 and 5.5. Peptides near the functional residue Val72, including residues 72–90, show pronounced acid-dependent increases in exchange, particularly at pH 5.5, consistent with enhanced local backbone dynamics and/or solvent exposure. In contrast, many longer peptides spanning residues 91–127 display comparatively modest pH-dependent changes, suggesting that this region remains relatively protected from acid-induced exchange. Peptides covering residues 126–139, near functional residues Arg130 and Arg132, and residues 140–159 show stronger acid sensitivity. Peptides spanning residues 158–169 also exhibit increased exchange under acidic conditions.

**Supplementary Figure 14.**
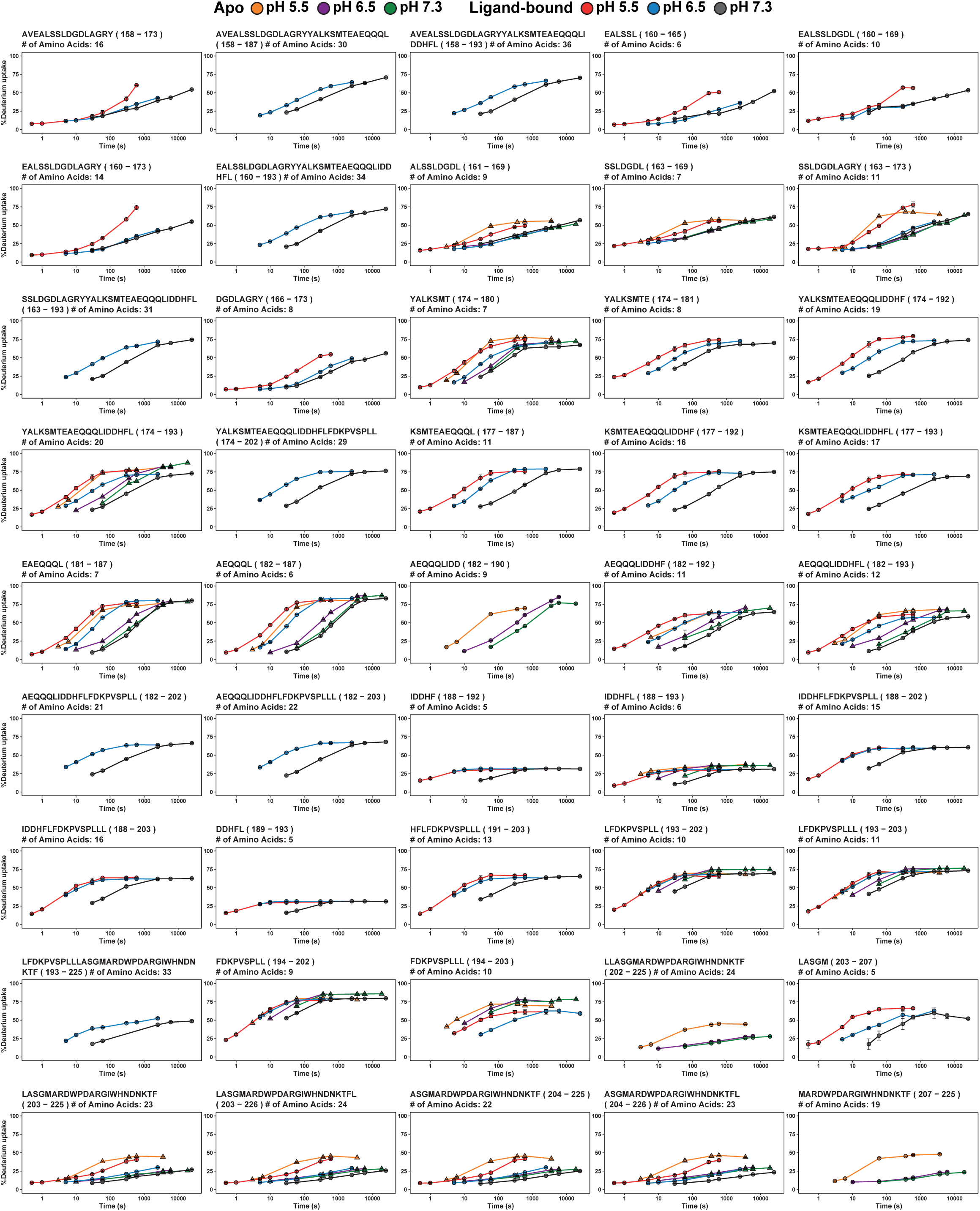
HDX uptake kinetics across the nucleotide-binding His191 and Ser199 phosphorylation region of the CK-BB concave surface. HDX uptake plots for overlapping peptides spanning residues 158–226. Peptides spanning the His191/Ser199 region, particularly residues 174–193 and 181–193, show pronounced pH-dependent exchange, with acidic pH increasing uptake and ligand binding producing a more graded pH response across the three conditions. In contrast, short peptides centered on residues 188–193 show comparatively limited exchange changes, suggesting local protection within the His191-containing segment. Peptides flanking this region, including residues 158–173 and 203–226, also display acid-enhanced exchange, indicating that pH-dependent backbone dynamics extend beyond the central His191/Ser199 site across the surrounding concave-surface region.

**Supplementary Figure 15.**
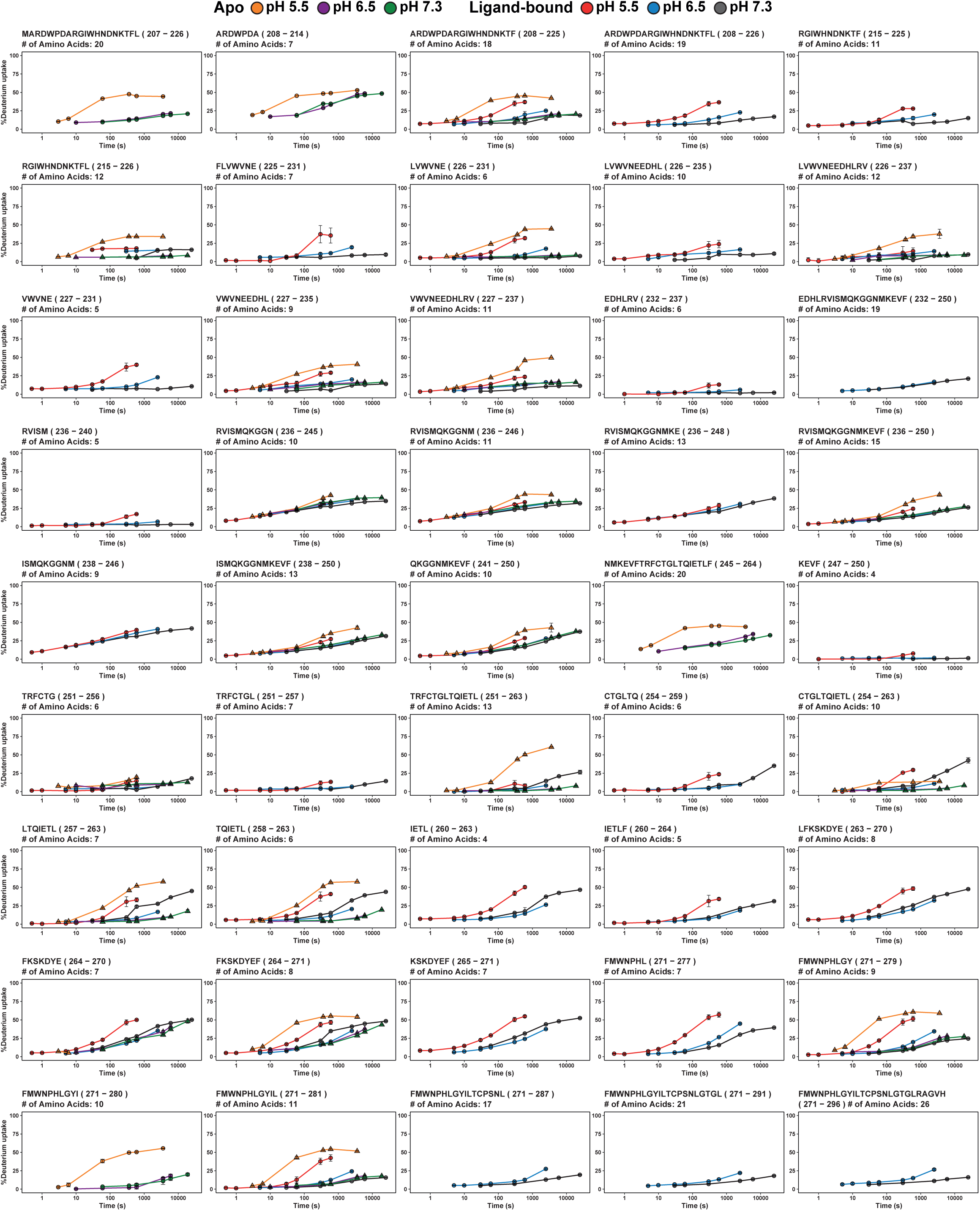
HDX uptake kinetics across catalytic and ligand-binding regions of CK-BB. HDX uptake plots for overlapping peptides spanning residues 207–296, shown as percentage deuterium uptake over exchange time for apo and ligand-bound CK-BB at pH 7.3, 6.5 and 5.5. This region includes catalytic and ligand-binding residues Glu232 and Arg236. Peptides spanning residues 207–226 show acid-enhanced exchange, particularly at pH 5.5, whereas several peptides across residues 226–250, including the Glu232/Arg236 region, remain comparatively protected with modest pH-dependent changes. In contrast, peptides spanning residues 251–281 display stronger acid-dependent increases in exchange, especially in the apo state and in selected ligand-bound traces, consistent with enhanced local backbone dynamics and/or solvent exposure.

**Supplementary Figure 16.**
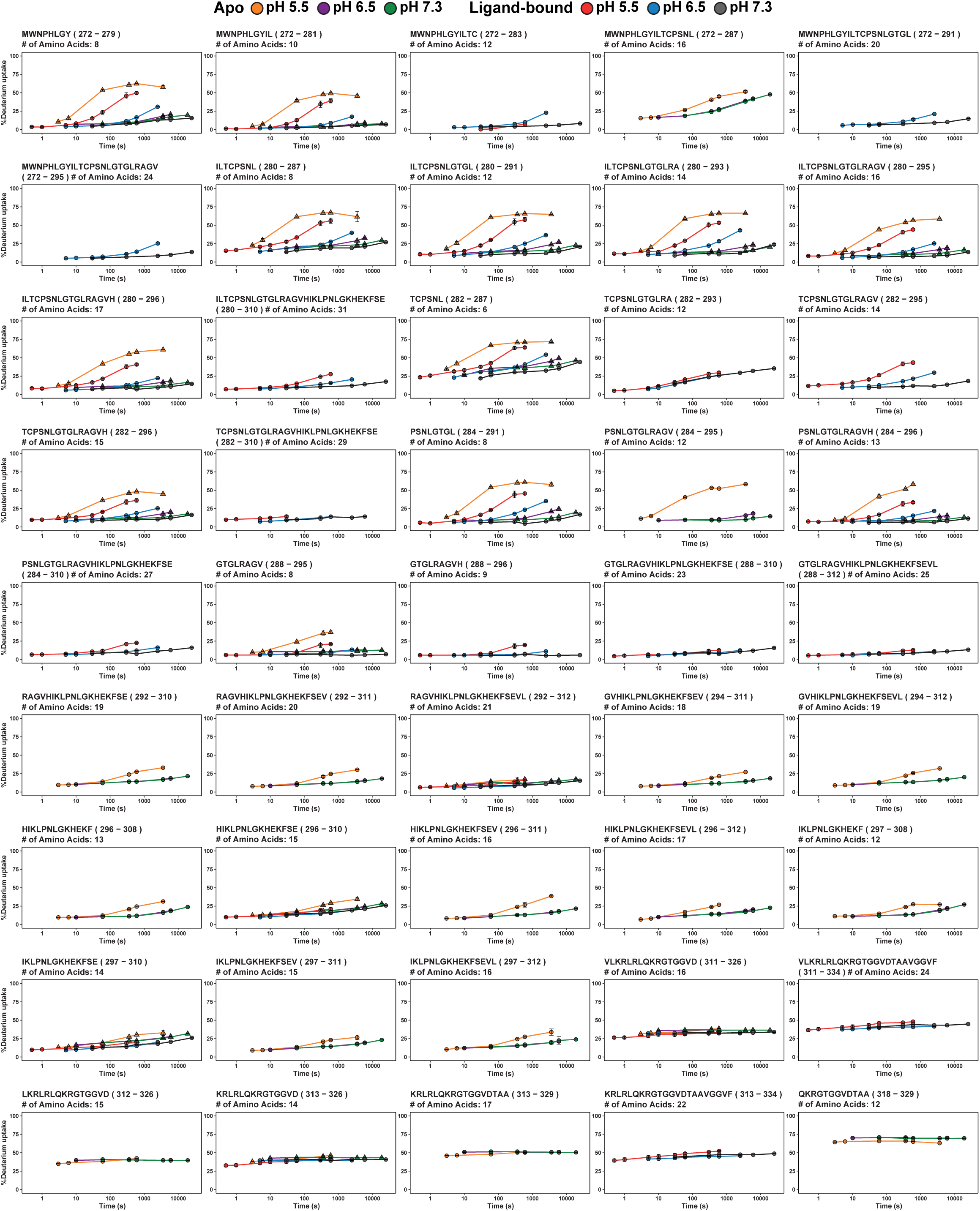
HDX uptake kinetics across the CK-BB CTD catalytic core. HDX uptake plots for overlapping peptides spanning residues 272–334. This region includes catalytic and ligand-binding residues Cys283, Arg292 and His296. Peptides spanning residues 272–296 show pronounced acid-dependent increases in exchange, particularly at pH 5.5 in the apo state, consistent with enhanced local backbone dynamics and/or solvent exposure around the Cys283/Arg292/His296 region. Ligand binding produces a more graded pH response across several of these peptides. In contrast, peptides spanning residues 292–312 show lower overall exchange and limited pH dependence, suggesting a comparatively protected CTD-core segment. Peptides spanning residues 311–334 are largely pH- and ligand-invariant, despite relatively high baseline exchange for some peptides, indicating constitutive exposure or flexibility rather than acid-specific remodeling.

**Supplementary Figure 17.**
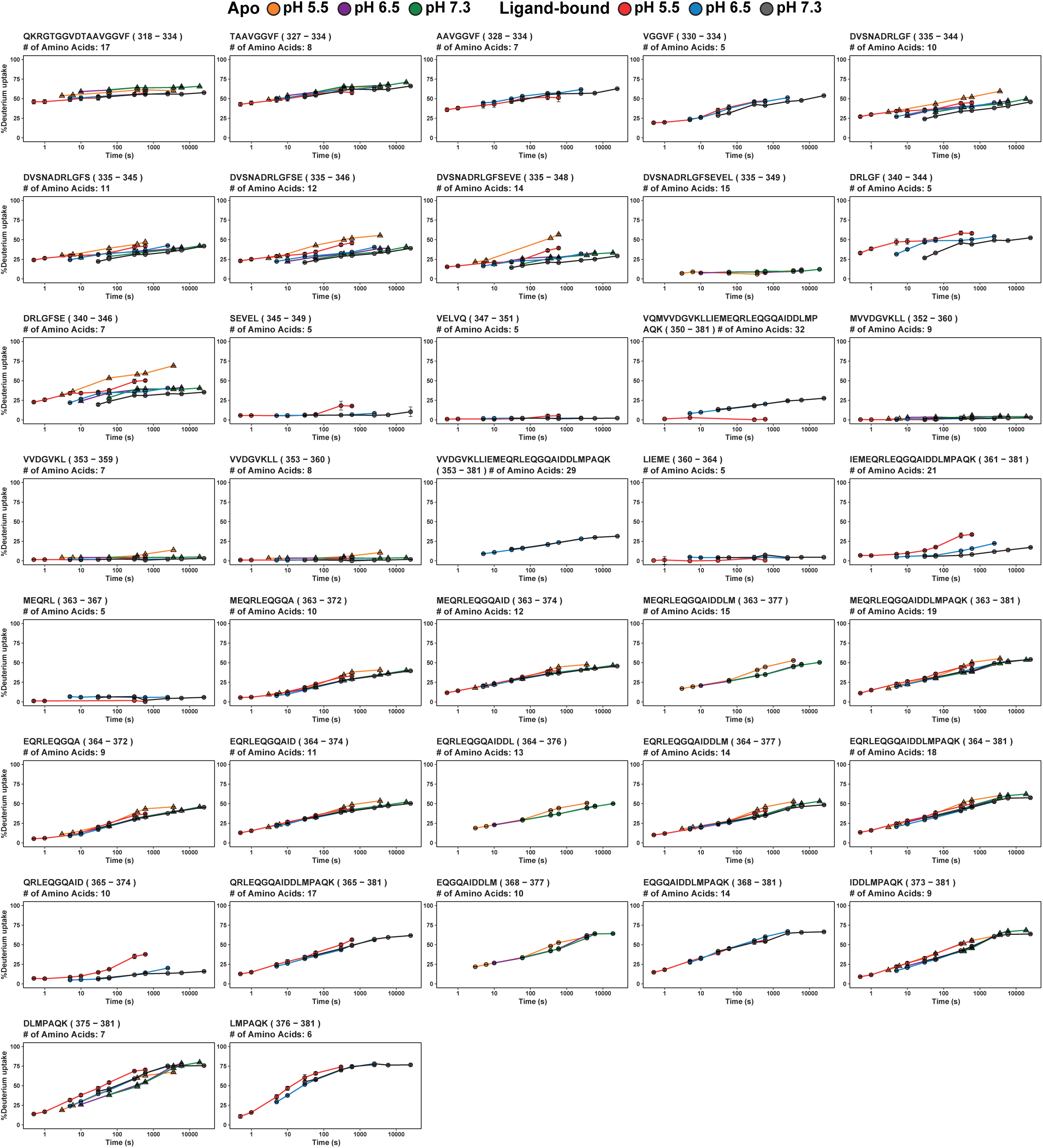
HDX uptake kinetics across the C-terminal region of the CK-BB CTD. HDX uptake plots for overlapping peptides spanning residues 318–381, shown as percentage deuterium uptake over exchange time for apo and ligand-bound CK-BB at pH 7.3, 6.5 and 5.5. This region includes functional residues Arg320, Val325 and Asp326. Peptides spanning residues 318–334 show relatively high baseline exchange with minimal pH- or ligand-dependent changes, suggesting constitutive solvent exposure or flexibility. Peptides within residues 335–348 display modest acid-dependent increases in exchange, most evident at pH 5.5, whereas short peptides around residues 345–351 show little exchange and limited condition dependence. In contrast, the distal C-terminal peptides spanning residues 363–381 exhibit gradual exchange over time with largely overlapping pH and ligand conditions, indicating a comparatively stable and condition-invariant C-terminal segment.

**Supplementary Table 1.** pH-adjusted HDX labeling times. HDX labeling times were adjusted using theoretical pH correction factors to compensate for slower intrinsic backbone amide exchange at lower pH. Relative to pH 7.3, exchange times were extended by factors of 6.3, 20.0 and 63.1 at pH 6.5, 6.0 and 5.5, respectively.

| pH | Time<br>(factor) | Experimental labeling times (s) |  |  |  |  |  |  |  |  |
| --- | --- | --- | --- | --- | --- | --- | --- | --- | --- | --- |
| 5.50 | 63.1 | 189 | 379 | 631 | 3786 | 22714 | 37857 | 227145 | 378574 | 1196737 |
| 6.00 | 20.0 | 60 | 120 | 200 | 1197 | 7183 | 11972 | 71829 | 119716 | 378441 |
| 6.50 | 6.3 | 19 | 38 | 63 | 379 | 2271 | 3786 | 22714 | 37857 | 119674 |
| 7.30 | 1.0 | 3 | 6 | 10 | 60 | 360 | 600 | 3600 | 6000 | 18967 |

## References

1. Wallimann, T. et al. Intracellular compartmentation, structure and function of creatine-kinase isoenzymes in tissues with high and fluctuating energy demands: the phosphocreatine circuit for cellular-energy homeostasis. Biochemical Journal 281, 21–40 (1992).

2. Dzeja, P. P. & Terzic, A. Phosphotransfer networks and cellular energetics. Journal of Experimental Biology 206, 2039–2047 (2003).

3. Wyss, M. & Kaddurah-Daouk, R. Creatine and creatinine metabolism. Physiological Reviews 80, 1107–1213 (2000).

4. Wallimann, T. & Hemmer, W. Creatine kinase in nonmuscle tissues and cells. Molecular and Cellular Biochemistry 133, 193–220 (1994).

5. Wallimann, T., Tokarska-Schlattner, M. & Schlattner, U. The creatine kinase system and pleiotropic effects of creatine. Amino Acids 40, 1271–1296 (2011).

6. Schlattner, U. et al. Cellular compartmentation of energy metabolism: creatine kinase microcompartments and recruitment of B-type creatine kinase to specific subcellular sites. Amino Acids 48, 1751–1774 (2016).

7. Jost, C. R. et al. Creatine kinase B-driven energy transfer in the brain is important for habituation and spatial learning behaviour, mossy fibre field size and determination of seizure susceptibility. European Journal of Neuroscience 15, 1692–1706 (2002).

8. Chang, E. J. et al. Brain-type creatine kinase has a crucial role in osteoclast-mediated bone resorption. Nature Medicine 14, 966–972 (2008).

9. Kuiper, J. W. P. et al. Creatine kinase-mediated ATP supply fuels actin-based events in phagocytosis. PLOS Biology 6, e51 (2008).

10. Kuiper, J. W. P. et al. Local ATP generation by brain-type creatine kinase (CK-B) facilitates cell motility. PLOS ONE 4, e5030 (2009).

11. Loo, J. M. et al. Extracellular metabolic energetics can promote cancer progression. Cell 160, 393–406 (2015).

12. Huddleston, H. G. et al. Clinical applications of microarray technology: creatine kinase B is an up-regulated gene in epithelial ovarian cancer and shows promise as a serum marker. Gynecologic Oncology 96, 77–83 (2005).

13. Tlili, M. et al. Creatine kinase B promotes non-small cell lung cancer survival and metastasis. Journal of Biological Chemistry 301, 110805 (2025).

14. Zweig, M. H. et al. Serum creatine-kinase isoenzyme-BB as an indicator of active metastatic disease. Clinical Chemistry 25, 1190–1191 (1979).

15. Ishiguro, Y. et al. The diagnostic and prognostic value of pretreatment serum creatine kinase-BB levels in patients with neuroblastoma. Cancer 65, 2014–2019 (1990).

16. Krutilina, R. I. et al. HIF-dependent CKB expression promotes breast cancer metastasis, whereas cyclocreatine therapy impairs cellular invasion and improves chemotherapy efficacy. Cancers 14, 27 (2022).

17. Li, X. H. et al. Knockdown of creatine kinase B inhibits ovarian cancer progression by decreasing glycolysis. International Journal of Biochemistry & Cell Biology 45, 979–986 (2013).

18. Papalazarou, V. et al. The creatine-phosphagen system is mechanoresponsive in pancreatic adenocarcinoma and fuels invasion and metastasis. Nature Metabolism 2, 62–80 (2020).

19. Kurth, I. et al. Therapeutic targeting of SLC6A8 creatine transporter suppresses colon cancer progression and modulates human creatine levels. Science Advances 7, eabi7511 (2021).

20. Darabedian, N. et al. Depletion of creatine phosphagen energetics with a covalent creatine kinase inhibitor. Nature Chemical Biology 19, 815–823 (2023).

21. Rashidi, A. et al. Myeloid cell-derived creatine in the hypoxic niche promotes glioblastoma growth. Cell Metabolism 36, 163–177 (2024).

22. Katz, J. L. et al. A covalent creatine kinase inhibitor ablates glioblastoma migration and sensitizes tumors to oxidative stress. Scientific Reports 14, 21959 (2024).

23. Wu, K. et al. Creatine kinase B suppresses ferroptosis by phosphorylating GPX4 through a moonlighting function. Nature Cell Biology 25, 714–725 (2023).

24. Pei, Y. F. et al. A current view on molecular mechanisms and machineries driving unconventional pathways of protein secretion. Current Opinion in Cell Biology 100, 102632 (2026).

25. Rabouille, C. Pathways of unconventional protein secretion. Trends in Cell Biology 27, 230–240 (2017).

26. Venter, G. et al. Submembranous recruitment of creatine kinase B supports formation of dynamic actin-based protrusions of macrophages and relies on its C-terminal flexible loop. European Journal of Cell Biology 94, 114–127 (2015).

27. Rojo, M. et al. Interaction of mitochondrial creatine kinase with model membranes: a monolayer study. FEBS Letters 281, 123–129 (1991).

28. Friedhoff, A. J. & Lerner, M. H. Creatine-kinase isoenzyme associated with synaptosomal membrane and synaptic vesicles. Life Sciences 20, 867–873 (1977).

29. Lim, L. et al. Neuron-specific enolase and creatine-phosphokinase are protein components of rat-brain synaptic plasma membranes. Journal of Neurochemistry 41, 1177–1182 (1983).

30. David, S., Shoemaker, M. & Haley, B. E. Abnormal properties of creatine kinase in Alzheimer’s disease brain: correlation of reduced enzyme activity and active site photolabeling with aberrant cytosol-membrane partitioning. Molecular Brain Research 54, 276–287 (1998).

31. Aksenov, M. et al. Oxidative modification of creatine kinase BB in Alzheimer’s disease brain. Journal of Neurochemistry 74, 2520–2527 (2000).

32. Bürklen, T. S. et al. The creatine kinase/creatine connection to Alzheimer’s disease: CK inactivation, APP-CK complexes, and focal creatine deposits. Journal of Biomedicine and Biotechnology 2006, 35936 (2006).

33. Rankin, E. B. & Giaccia, A. J. Hypoxic control of metastasis. Science 352, 175–180 (2016).

34. Gatenby, R. A. & Gillies, R. J. Why do cancers have high aerobic glycolysis? Nature Reviews Cancer 4, 891–899 (2004).

35. Estrella, V. et al. Acidity generated by the tumor microenvironment drives local invasion. Cancer Research 73, 1524–1535 (2013).

36. Monastyrskaya, K. et al. Annexins sense changes in intracellular pH during hypoxia. Biochemical Journal 409, 65–75 (2008).

37. Tyrtyshnaia, A. A. et al. Acute neuroinflammation provokes intracellular acidification in mouse hippocampus. Journal of Neuroinflammation 13, 283 (2016).

38. Prasad, H. & Rao, R. Amyloid clearance defect in ApoE4 astrocytes is reversed by epigenetic correction of endosomal pH. Proceedings of the National Academy of Sciences of the United States of America 115, E6640–E6649 (2018).

39. Eder, M. et al. Crystal structure of brain-type creatine kinase at 1.41 Å resolution. Protein Science 8, 2258–2269 (1999).

40. Lumpkin, R. J. et al. Structure and dynamics of the ASB9 CUL-RING E3 ligase. Nature Communications 11, 2866 (2020).

41. Bong, S. M. et al. Structural studies of human brain-type creatine kinase complexed with the ADP-Mg-NO-creatine transition-state analogue complex. FEBS Letters 582, 3959–3965 (2008).

42. Forstner, M. et al. The active site histidines of creatine kinase: a critical role of His61 situated on a flexible loop. Protein Science 6, 331–339 (1997).

43. Wood, T. D. et al. Creatine kinase: essential arginine residues at the nucleotide binding site identified by chemical modification and high-resolution tandem mass spectrometry. Proceedings of the National Academy of Sciences of the United States of America 95, 3362–3365 (1998).

44. Lahiri, S. D. et al. The 2.1 Å structure of creatine kinase complexed with the ADP-Mg-NO-creatine transition-state analogue complex. Biochemistry 41, 13861–13867 (2002).

45. Wang, P. F. et al. Loop movement and catalysis in creatine kinase. IUBMB Life 57, 355–362 (2005).

46. Polino, A. J. et al. Disrupting actin filaments enhances glucose-stimulated insulin secretion independent of the cortical actin cytoskeleton. Journal of Biological Chemistry 299, 105334 (2023).

47. Schiemann, O. et al. Benchmark test and guidelines for DEER/PELDOR experiments on nitroxide-labeled biomolecules. Journal of the American Chemical Society 143, 17875–17890 (2021).

48. Dastvan, R. et al. Proton-driven alternating access in a spinster lipid transporter. Nature Communications 13, 5161 (2022).

49. Verhalen, B. et al. Energy transduction and alternating access of the mammalian ABC transporter P-glycoprotein. Nature 543, 738–741 (2017).

50. Dastvan, R. & Stoll, S. Recent advances in quantifying protein conformational ensembles with dipolar EPR spectroscopy. Current Opinion in Structural Biology 94, 103139 (2025).

51. Assafa, T. E. et al. Light-driven domain mechanics of a minimal phytochrome photosensory module studied by EPR. Structure 26, 1534–1543.e4 (2018).

52. Dikiy, I. et al. Insights into histidine kinase activation mechanisms from the monomeric blue light sensor EL346. Proceedings of the National Academy of Sciences of the United States of America 116, 4963–4972 (2019).

53. Wu, T. Q. et al. Modeling protein conformational ensembles by guiding AlphaFold2 with double electron-electron resonance distance distributions. Nature Communications 16, 7107 (2025).

54. Dastvan, R. et al. Protonation-dependent conformational dynamics of the multidrug transporter EmrE. Proceedings of the National Academy of Sciences of the United States of America 113, 1220–1225 (2016).

55. Jumper, J. et al. Highly accurate protein structure prediction with AlphaFold. Nature 596, 583–589 (2021).

56. Abramson, J. et al. Accurate structure prediction of biomolecular interactions with AlphaFold 3. Nature 630, 493–500 (2024).

57. McLaughlin, N. K. et al. Kingfisher: an open-sourced web-based platform for the analysis of hydrogen exchange mass spectrometry data. Protein Science 34, e70096 (2025).

58. Li, J. et al. Hydrogen–deuterium exchange and mass spectrometry reveal the pH-dependent conformational changes of diphtheria toxin T domain. Biochemistry 53, 6849–6856 (2014).

59. Rodnin, M. V. et al. The pH-dependent trigger in diphtheria toxin T domain comes with a safety latch. Biophysical Journal 111, 1946–1953 (2016).

60. Johnson, B. et al. Allosteric coupling of CARMIL and V-1 binding to capping protein revealed by hydrogen–deuterium exchange. Cell Reports 23, 2795–2804 (2018).

61. Vadas, O. et al. Using hydrogen–deuterium exchange mass spectrometry to examine protein-membrane interactions. Methods in Enzymology 583, 143–172 (2017).

62. Lane, B. J. et al. HDX-guided EPR spectroscopy to interrogate membrane protein dynamics. STAR Protocols 3, 101562 (2022).

63. Moradi, M. & Tajkhorshid, E. Mechanistic picture for conformational transition of a membrane transporter at atomic resolution. Proceedings of the National Academy of Sciences of the United States of America 110, 18916–18921 (2013).

64. Moradi, M. & Tajkhorshid, E. Computational recipe for efficient description of large-scale conformational changes in biomolecular systems. Journal of Chemical Theory and Computation 10, 2866–2880 (2014).

65. Moradi, M., Enkavi, G. & Tajkhorshid, E. Atomic-level characterization of transport cycle thermodynamics in the glycerol-3-phosphate:phosphate antiporter. Nature Communications 6, 8393 (2015).

66. Govind Kumar, V., et al. Binding affinity estimation from restrained umbrella sampling simulations. Nature Computational Science 3, 59–70 (2023).

67. Badiee, S. A., Govind Kumar, V. & Moradi, M. Molecular dynamics investigation of the influenza hemagglutinin conformational changes in acidic pH. Journal of Physical Chemistry B 128, 11151–11163 (2024).

68. Zhao, T. J. et al. Impact of intra-subunit domain-domain interactions on creatine kinase activity and stability. FEBS Letters 580, 3835–3840 (2006).

69. Zhao, T. J. et al. The generation of the oxidized form of creatine kinase is a negative regulation on muscle creatine kinase. Journal of Biological Chemistry 282, 12022–12029 (2007).

70. Furter, R., Furtergraves, E. M. & Wallimann, T. Creatine kinase: the reactive cysteine is required for synergism but is nonessential for catalysis. Biochemistry 32, 7022–7029 (1993).

71. Zeno, W. F. et al. Synergy between intrinsically disordered domains and structured proteins amplifies membrane curvature sensing. Nature Communications 9, 4152 (2018).

72. Johnson, D. H. et al. Lipid packing defects are necessary and sufficient for membrane binding of α-synuclein. Communications Biology 8, 1179 (2025).

73. Ponticos, M. et al. Dual regulation of the AMP-activated protein kinase provides a novel mechanism for the control of creatine kinase in skeletal muscle. EMBO Journal 17, 1688–1699 (1998).

74. James, E. I. et al. Advances in hydrogen/deuterium exchange mass spectrometry and the pursuit of challenging biological systems. Chemical Reviews 122, 7562–7623 (2022).

75. Hamuro, Y. Interpretation of hydrogen/deuterium exchange mass spectrometry. Journal of the American Society for Mass Spectrometry 35, 819–828 (2024).

76. Hamuro, Y. Quantitative hydrogen/deuterium exchange mass spectrometry. Journal of the American Society for Mass Spectrometry 32, 2711–2727 (2021).

77. Englander, J. J. et al. Protein structure change studied by hydrogen–deuterium exchange, functional labeling, and mass spectrometry. Proceedings of the National Academy of Sciences of the United States of America 100, 7057–7062 (2003).

78. Gonzalez, F. A., Raden, D. L. & Davis, R. J. Identification of substrate recognition determinants for human ERK1 and ERK2 protein kinases. Journal of Biological Chemistry 266, 22159–22163 (1991).

79. Michl, J., Park, K. C. & Swietach, P. Evidence-based guidelines for controlling pH in mammalian live-cell culture systems. Communications Biology 2, 144 (2019).

80. Kou, O. H. et al. Membrane phase, charge, and curvature regulate α-synuclein binding dynamics. Langmuir 42, 25681–25696 (2026).

81. Kou, O. H., Kim, B. H., Johnson, D. H. & Zeno, W. F. Cholesterol differentially regulates α-synuclein binding across membrane packing regimes. Biophysical Reports 6, 100282 (2026).

82. Zeno, W. F. et al. Molecular mechanisms of membrane curvature sensing by a disordered protein. Journal of the American Chemical Society 141, 10361–10371 (2019).

83. Jeschke, G. DEER distance measurements on proteins. Annual Review of Physical Chemistry 63, 419–446 (2012).

84. Hustedt, E. J., Stein, R. A. & McHaourab, H. S. Protein functional dynamics from the rigorous global analysis of DEER data: conditions, components, and conformations. Journal of General Physiology 153, e201711954 (2021).

85. Polyhach, Y., Bordignon, E. & Jeschke, G. Rotamer libraries of spin-labelled cysteines for protein studies. Physical Chemistry Chemical Physics 13, 2356–2366 (2011).

86. Mirdita, M. et al. ColabFold: making protein folding accessible to all. Nature Methods 19, 679–682 (2022).

87. Wu, T. Q. et al. Analysis of several key factors influencing deep learning-based inter-residue contact prediction. Bioinformatics 36, 1091–1098 (2020).

88. Sali, A. & Blundell, T. L. Comparative protein modeling by satisfaction of spatial restraints. Journal of Molecular Biology 234, 779–815 (1993).

89. Tamara, S., den Boer, M. A. & Heck, A. J. R. High-resolution native mass spectrometry. Chemical Reviews 122, 7269–7326 (2022).

90. Karch, K. R. et al. Native mass spectrometry: recent progress and remaining challenges. Annual Review of Biophysics 51, 157–179 (2022).

91. Olsson, M. H. M. et al. PROPKA3: consistent treatment of internal and surface residues in empirical pKa predictions. Journal of Chemical Theory and Computation 7, 525–537 (2011).

92. Lee, J. et al. CHARMM-GUI input generator for NAMD, GROMACS, AMBER, OpenMM and CHARMM/OpenMM simulations using the CHARMM36 additive force field. Biophysical Journal 110, 641a (2016).

93. Jorgensen, W. L. et al. Comparison of simple potential functions for simulating liquid water. Journal of Chemical Physics 79, 926–935 (1983).

94. Huang, J. et al. CHARMM36m: an improved force field for folded and intrinsically disordered proteins. Nature Methods 14, 71–73 (2017).

95. Phillips, J. C. et al. Scalable molecular dynamics on CPU and GPU architectures with NAMD. Journal of Chemical Physics 153, 044130 (2020).

96. Feller, S. E. et al. Constant-pressure molecular-dynamics simulation: the Langevin piston method. Journal of Chemical Physics 103, 4613–4621 (1995).

97. Shaw, D. E. et al. Anton 3: twenty microseconds of molecular dynamics simulation before lunch. In SC ’21: Proceedings of the International Conference for High Performance Computing, Networking, Storage and Analysis 1–11 (Association for Computing Machinery, 2021).

98. Humphrey, W., Dalke, A. & Schulten, K. VMD: visual molecular dynamics. Journal of Molecular Graphics and Modelling 14, 33–38 (1996).

## References

1. Momin, N. et al. Designing lipids for selective partitioning into liquid ordered membrane domains. Soft Matter 11, 3241–3250 (2015).

2. Johnson, D. H. et al. Lipid packing defects are necessary and sufficient for membrane binding of α-synuclein. Communications Biology 8, 1179 (2025).

3. Kou, O. H. et al. Membrane phase, charge, and curvature regulate α-synuclein binding dynamics. Langmuir 42, 25681–25696 (2026).

4. Kou, O. H., Kim, B. H., Johnson, D. H. & Zeno, W. F. Cholesterol differentially regulates α-synuclein binding across membrane packing regimes. Biophysical Reports 6, 100282 (2026).

5. Zeno, W. F. et al. Synergy between intrinsically disordered domains and structured proteins amplifies membrane curvature sensing. Nature Communications 9, 4152 (2018).

6. Zeno, W. F. et al. Molecular mechanisms of membrane curvature sensing by a disordered protein. Journal of the American Chemical Society 141, 10361–10371 (2019).

7. McLaughlin, N. K. et al. Kingfisher: An open-sourced web-based platform for the analysis of hydrogen exchange mass spectrometry data. Protein Science 34, e70096 (2025).

8. Hageman, T. S. & Weis, D. D. Reliable identification of significant differences in differential hydrogen exchange-mass spectrometry measurements using a hybrid significance testing approach. Analytical Chemistry 91, 8008–8016 (2019).

9. Lau, A. M., Claesen, J., Hansen, K. & Politis, A. Deuteros 2.0: peptide-level significance testing of data from hydrogen deuterium exchange mass spectrometry. Bioinformatics 37, 270–272 (2021).

